# *Mycobacterium tuberculosis* FtsB and PerM interact via a C-terminal helix in FtsB to modulate cell division

**DOI:** 10.1101/2024.03.11.584518

**Authors:** Joaõ Ramalheira Ferreira, Ruilan Xu, Zach Hensel

## Abstract

Latent infection by *Mycobacterium tuberculosis* (Mtb) impedes effective tuberculosis therapy and eradication. The protein PerM is essential for chronic Mtb infections in mice and acts via the divisome protein FtsB to modulate cell division. Using transgenic co-expression in *Escherichia coli*, we studied the Mtb PerM-FtsB interaction in isolation from other Mtb proteins, engineering PerM to enhance expression in the *E. coli* membrane. We confirmed the reported instability of Mtb FtsB, and we linked FtsB instability to a segment of FtsB predicted to bind cell-division proteins FtsL and FtsQ. Using fluorescence microscopy, we found that PerM stability hinged on its interaction with a C-terminal helix in FtsB. Molecular dynamics results supported the observation that FtsB stabilized PerM, and suggested that interactions at the PerM-FtsB interface differ from our initial structure prediction in a way that is consistent with PerM sequence conservation. Though narrowly conserved, the PerM-FtsB interaction emerges as a potential target for therapy targeting persistent infections by disrupting regulation of cell division. Integrating protein structure prediction, molecular dynamics and single-molecule microscopy, our approach is primed to screen potential inhibitors of the PerM-FtsB interaction and can be straightforwardly adapted to explore other putative interactions.

**Importance:** Our research reveals significant insights into the dynamic interaction between the proteins PerM and FtsB within *Mycobacterium tuberculosis*, contributing to our understanding of bacterial cell division mechanisms crucial for infection persistence. By combining innovative fluorescence microscopy and molecular dynamics, we established that the stability of these proteins is interdependent; molecular dynamics placing PerM-FtsB in the context of the mycobacterial divisome show how disrupting PerM-FtsB interaction can plausibly impact bacterial cell wall synthesis. These findings highlight the PerM-FtsB interface as a promising target for novel therapeutics aimed at combating persistent bacterial infections. Importantly, our approach can be adapted for similar studies in other bacterial systems, suggesting broad implications for microbial biology and antibiotic development.

## Introduction

*Mycobacterium tuberculosis* (Mtb) is a major contributor to preventable deaths by infectious disease and has infected approximately a quarter of the global population [1]. Most infections progress to latent tuberculosis infection (LTBI), characterized by an immunological response to Mtb antigens without clinical signs of active TB [2]. Preventing establishment and reactivation of LTBI is a growing priority for TB elimination strategies, although uncertainties continue to limit measurements of the relative burdens of new TB infections and LTBI reactivation [3]. Protocols that reduce the duration and adverse effects of treatment can reduce LTBI burden by improving completion rates for preventative treatment [4]. Furthermore, imperfect treatment over long periods can contribute to acquisition of drug resistance [5]. Consequently, it is imperative to better understand how Mtb infections persist to establish LTBI in order to inform strategies for novel LTBI treatments with higher efficacy, shorter durations, and reduced risk of drug resistance.

A potential strategy to prevent establishment and reactivation of LTBI is to characterize and target mechanisms through which Mtb persists through host-induced stress [6]. The actinomycete protein PerM was one of 21 genes identified in a screen of transposon Mtb mutants with attenuated growth at pH 4.5 in media containing Tween 80, with most of the identified genes being associated with cell wall synthesis [7]. A subsequent study focused on PerM, finding that Mtb PerM knockout reduced growth during chronic mouse infection and increased *β*-lactam antibiotic susceptibility. Localization of fluorescently tagged PerM to dividing septa suggested a connection to cell division [8]. Further work demonstrated that PerM associates with the Mtb divisome, a protein complex orchestrating cell wall remodeling during division, and that PerM depletion can be complemented by overexpression of the divisome component FtsB [9]. PerM was also recently identified by transposon sequencing to play an even more significant role in infecting mice in a background with weakened adaptive immune response [10].

Recently, a covalent inhibitor of divisome formation based upon the structure of FtsB-FtsQ was developed and found to be active against drug-resistant *E. coli* in an animal infection model [11]. The Mtb proteome includes homologs of the five core *E. coli* divisome proteins FtsQ, FtsL, FtsB, FtsW, and FtsI [12], suggesting that a similar approach can target protein-protein interactions in the Mtb divisome. However, it is unknown whether PerM directly interacts with FtsB [9] and, if it does, whether PerM has a regulatory role beyond impacting FtsB stability. PerM lacks known orthologs outside of actinomycetes [8], so PerM interactions could potentially be targeted by specific therapies. However, no experimental structural data have been published on the PerM-FtsB interaction or on PerM alone to guide experimental design. Expression and purification of Mtb PerM and FtsB for structural study is likely to pose difficulties given that FtsB stability depends upon PerM co-expression in Mtb and *Mycolicibacterium smegmatis* [9]. Furthermore, proteins such as PerM with a high number of transmembrane helices pose challenges for recombinant expression [13, 14].

Recent insights into the molecular structure and regulatory mechanisms of the gram-negative divisome have been enabled by the confluence of protein structure prediction, cryogenic electron microscopy (Cryo-EM), and molecular dynamics (MD) simulations. The extension from prediction of monomers to protein complexes [15, 16] facilitated rapid structure prediction of the core *E. coli* divisome [17, 18]. Analysis of a Cryo-EM structure of the *Pseudomonas aeruginosa* divisome was largely consistent with predicted protein-protein interfaces, but also revealed a global conformational change absent in structure predictions [19]. All-atom MD simulations identified a similar conformational change within 1 µs, and further predicted interactions between the core divisome and *E. coli* FtsN [20]. However, limitations of this MD approach, such as uncertainty in structure predictions and limited MD timescales, call for experimental validation. Importantly, *in silico* predictions for FtsN were corroborated by experimental results in living cells [21].

In this work, we employed a combination of structure prediction, MD, and fluorescence microscopy to probe the predicted interaction between Mtb PerM and FtsB proteins. In our approach, Mtb PerM and FtsB were expressed *E. coli* in order to directly attribute observations to changes in the Mtb proteins. First, we investigated conservation and dynamics at the predicted PerM-FtsB interface, identifying roles for conserved residues that are absent in structure predictions. Second, we showed that FtsB instability (when expressed in *E. coli*) depends on a region predicted to bind FtsL and FtsQ, and that PerM expression in the *E. coli* membrane can be enhanced by strategically modifying its N-terminal signal sequence. This enabled the quantification of the PerM-FtsB interaction via fluorescence correlation and single-molecule tracking. We found PerM expression in the *E. coli* membrane to depend on the FtsB sequence predicted to mediate PerM-FtsB interaction. Lastly, we investigated MD simulations of Mtb divisome complexes suggested PerM regulatory complexity beyond FtsB stabilization.

## Results

### Structure prediction and molecular dynamics of PerM-FtsB interaction

In preliminary structure predictions, we identified a high-confidence predicted interaction between Mtb PerM and FtsB (*pDockQ* ≈ 0.6 [22]). The predicted structure of PerM was largely consistent with previous predicted topology [8] except that two regions with hydrophobic residues in predicted transmembrane helices did not span the membrane in structure predictions (Fig. 1A). The topology remains N-in, C-in, with the predicted PerM structure consisting of two halves, each with the same topology (four transmembrane helices with a buried extracellular loop between the first two). Preliminary molecular dynamics (MD) simulations showed that the predicted PerM-FtsB interaction persisted on the microsecond timescale. To test stability of the structure of the predicted complex further, we carried out a three-stage MD protocol with 100 ns of equilibrium MD followed by 500 ns of accelerated molecular dynamics (aMD), and a final 500 ns of equilibrium MD. Terminal residues lacking high local prediction confidence (*pLDDT* < 50) were omitted in constructing the simulation system of FtsB^61–204^ and PerM^11–390^. Fig. 1A shows how the predicted extended structure of FtsB collapsed in the absence of interactions with other divisome components [17–20]. Interaction between transmembrane helices of PerM and FtsB was not predicted with high confidence and also did not exhibit persistent, specific interactions in MD. Conversely, the predicted interaction between FtsB and a specific pocket in the periplasmic face of PerM was maintained throughout the aMD protocol and in a simulation replicate.

**Fig. 1.**
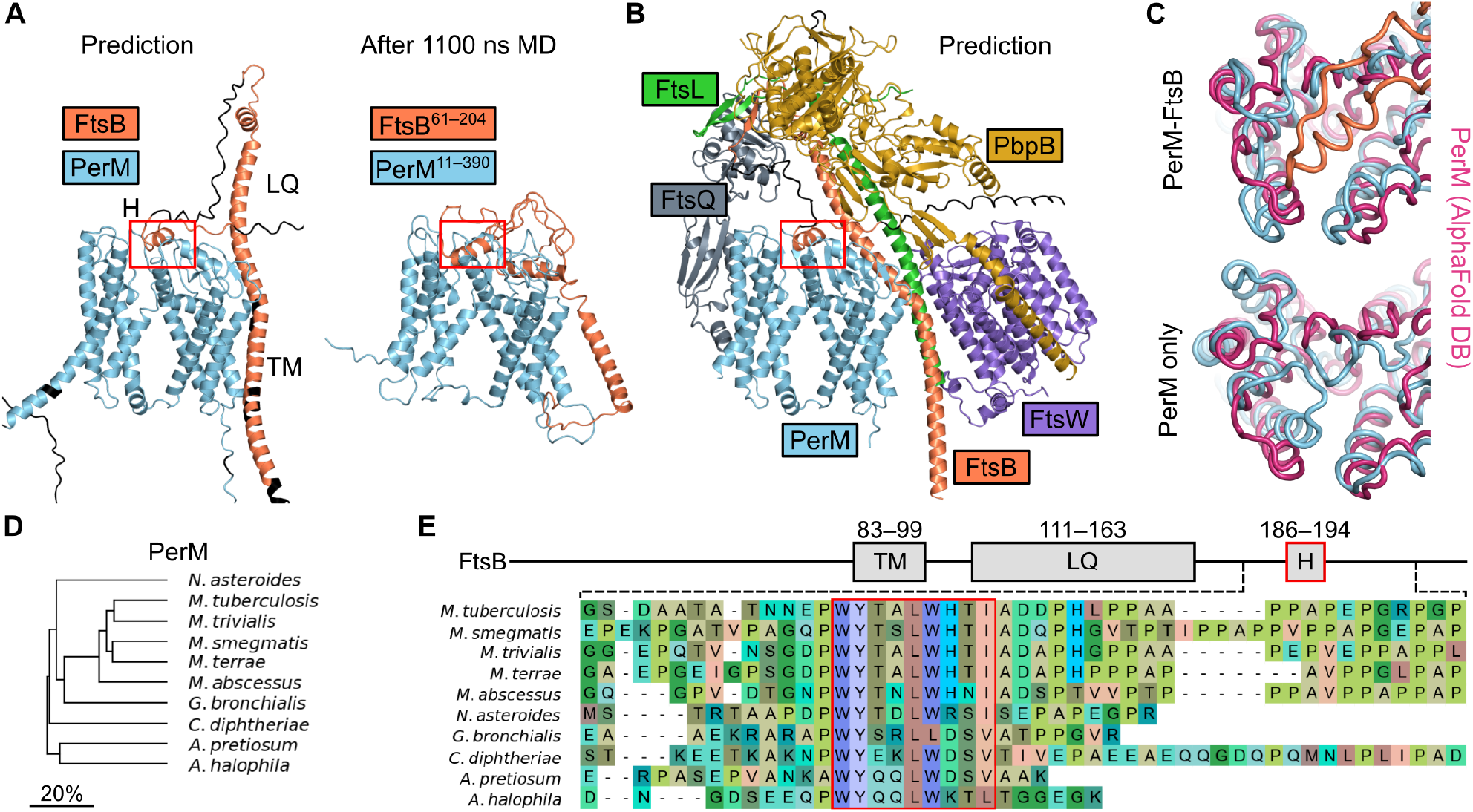
A conserved, predicted Mtb PerM-FtsB interaction stabilizes PerM in MD. (**A**) Left: predicted PerM-FtsB complex in which an *α* helix, FtsB^H^, interacts with the periplasmic face of PerM. Transmembrane helix FtsB™ and the region interacting with FtsL and FtsQ, FtsB^LQ^, are indicated. Residues with *pLDDT* < 50 colored black. Right: final conformer following 1.1 µs MD; PerM-FtsB^H^ interact persists, FtsB^LQ^ collapses, and FtsB™ moves away from PerM. (**B**) Prediction for the Mtb divisome with inclusion of PerM. Terminal regions with *pLDDT* < 50 are omitted except for the FtsB C-terminus. All complexes in **A** and **B** are aligned by PerM C*α* atoms; the red box is placed at the same position relative to PerM for comparison of FtsB^H^ movement. (**C**) Final MD conformers for PerM-FtsB and for PerM alone are aligned by PerM C*α* atoms and compared to a PerM prediction (AlphaFold DB P9WKN3-F1-model v4). (**D**) Phylogenetic tree for actinomycete species with predicted PerM-FtsB interactions. Branch length scaled by divergence in amino acid identity in pairwise sequence alignments of PerM. (**E**) Top: Diagram of Mtb FtsB defining the FtsB™, FtsB^LQ^, and FtsB^H^ regions. Bottom: Multiple sequence alignment of a region near the C-terminus of FtsB for actinomycete species illustrates conservation of residues in FtsB^H^ and diversity in FtsB C-termini.

We also found that a prediction of the core Mtb divisome with the addition of PerM was consistent with PerM interacting with the divisome without obviously disrupting interactions between core divisome components (Fig. 1B). Within this complex, PerM was only predicted to interact with FtsB. Based on these predictions, we identified three regions of interest in Mtb FtsB: the predicted transmembrane helix, FtsB™ (FtsB^83–99^), the region predicted to interact with FtsL and FtsQ, FtsB^LQ^ (FtsB^111–163^), and a span of hydrophobic residues forming an *α* helix predicted to interact with PerM, FtsB^H^ (FtsB^186–194^). Although C-terminal residues following FtsB^H^ were predicted to thread between the PbpB anchor and head domains, there were no high-confidence predictions for this region.

In order to see the impact of FtsB on PerM, we replicated the three-stage MD protocol using PerM alone. Final conformers for PerM-FtsB and PerM simulations are shown in Fig. 1C and compared to a PerM structure prediction from the AlphaFold Protein Structure Database [23]. Surprisingly, we found that the final conformer in the monomeric PerM simulation differed from the predicted PerM structure (*RMSD* = 3.47 °A) to a greater extent than final conformers in PerM-FtsB simulation replicates (2.50 °A and 2.21 °A). The largest structural changes were seen in the predicted FtsB^H^ binding pocket.

Since our structure predictions were informed by information encoded in multiple sequence alignments, we searched for and identified PerM orthologs in actinomycete species (Fig. 1D). For each, we also identified the corresponding FtsB sequence and predicted the structures of PerM-FtsB complexes. In every species tested there was a predicted interaction between PerM and FtsB^H^ with the same orientation of FtsB^H^ on the periplasmic face of PerM (Supplementary Fig. S1). Further, there is little difference in the lengths of regions linking FtsB^LQ^ and FtsB^H^. PerM-FtsB complexes differed, however, in the predicted position of FtsB™ relative to PerM. We constructed a multiple sequence alignment of FtsB to investigate the basis for conserved, predicted PerM-FtsB interaction and found that FtsB^H^ was highly conserved (Fig. 1E) with near perfect conservation of hydrophobic residues predicted to interact with PerM (W186, Y187, L190, and W191).

Next, we investigated the structural basis for specific PerM-FtsB interaction. Fig. 2A illustrates how the FtsB binding pocket of PerM changed during MD, focusing on hydrogen bonds formed during MD that were not observed in structure predictions. These include a sidechain-sidechain hydrogen bond with PerM Y239, which is perfectly conserved in PerM for the species shown in Fig. 1D. We observed that a large, hydrophobic binding pocket tightened around FtsB^H^, and around FtsB W186 in particular. Residues in the W186 binding pocket include PerM G228, which was conserved in mycobacterial sequences in species that we analyzed.

**Fig. 2.**
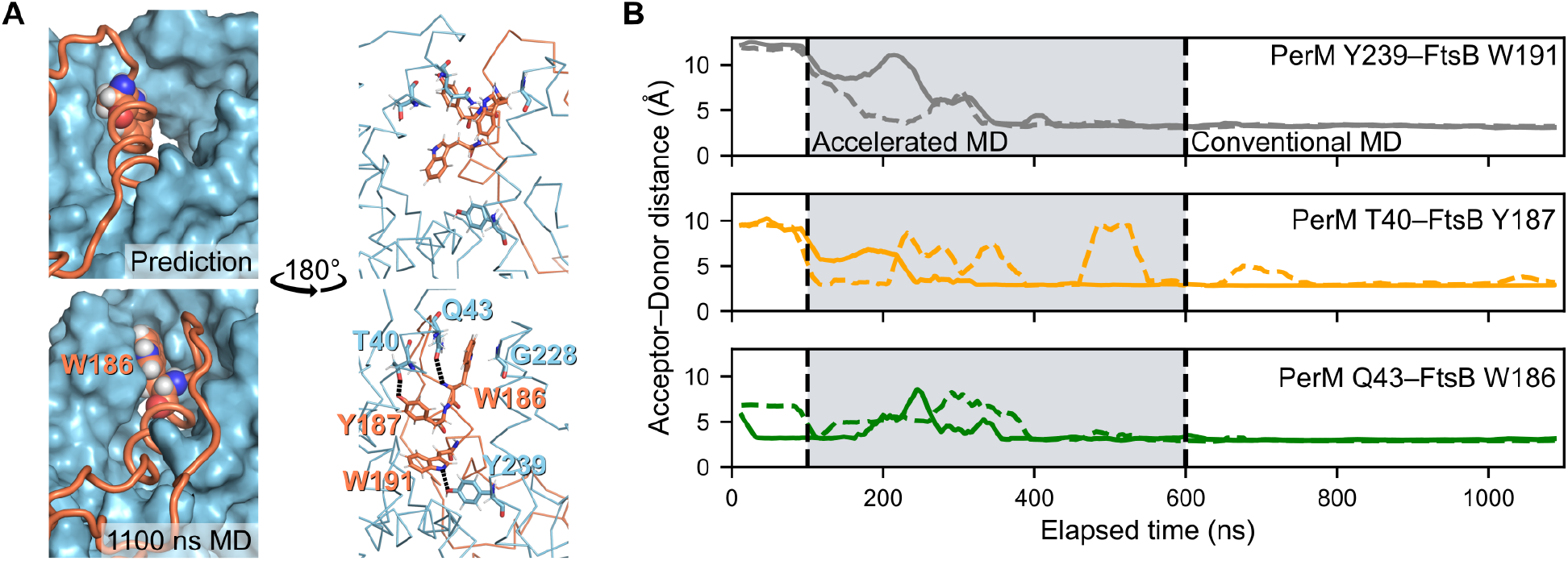
Reproducible dynamics at the PerM-FtsB binding interface. (**A**) The shape of the FtsB binding pocket for PerM is compared between the predicted interface structure (top) and the final conformer following 1.1 µs MD (bottom). Left: a large predicted periplasmic binding pocket in PerM (top left) evolves to tightly bind FtsB W186 (bottom left). Right: Following MD (bottom right), FtsB W186 interacts with conserved PerM residue G228 and hydrogen bonds are formed between conserved FtsB residues and PerM residues including highly conserved Y239. The same region is shown rotated by 180°. (**B**) Interactions at the PerM-FtsB interface appeared in two independent aMD simulations. Solid and dashed lines show acceptor-donor distance (25 ns moving average; two MD replicates) for hydrogen bonds at the PerM-FtsB interface that are absent in the predicted structure.

Fig. 2B shows that hydrogen bonding interactions at the PerM-FtsB binding interface were reproduced in a replicate simulation. However, we observed that the *α*-helical secondary structure of FtsB^H^ was partially lost in the second simulation (Supplementary Fig. S2A). To further assess reproducibility, we conducted two additional replicate aMD simulations (differing only in using shorter, 50 ns final steps) that also demonstrated loss of *α*-helical dihedral angles in FtsB^H^ and, further, failed to reproduce the interactions shown in Fig. 2A (Supplementary Fig. S2B). We suspected that a reduced dihedral boost in aMD would maintain secondary structure while still more rapidly exploring conformations at the PerM-FtsB interface than conventional MD. Two final simulations with reduced dihedral boost confirmed this hypothesis, maintaining *α*-helical dihedral angles and more frequently exhibiting the hydrogen bonds observed in our full-length simulations (Supplementary Fig. S2C).

### Transgenic expression of fluorescent FtsB and PerM constructs in *E. coli*

Despite the apparent specificity and stability of the predicted PerM-FtsB interface in MD, we wanted to confirm that this interaction does not depend on other Mtb components to be stable on timescales inaccessible by MD and, if so, to investigate whether FtsB^H^ mediated the interaction. Rather than reconstitute a minimal system *in vitro*, we aimed to investigate the bimolecular interaction in *E. coli* cells using Mtb FtsB and PerM fused to fluorescent proteins. We chose the fluorescent proteins mNeonGreen [24] and mTurquoise2 [25] since these bright, fast-maturing fluorescent proteins can be spectrally separated with appropriate filter sets.

We constructed variants of FtsB with an N-terminal fusion of mTurquoise2 (mTq2-FtsB, mTq2-FtsB^ΔLQ^, and mTq2-FtsB^ΔLQΔH^; Fig. 3A). Deletion of the FtsB^LQ^ region was based upon the observation of hydrophobic interactions of Mtb FtsB with FtsL and FtsQ in the predicted Mtb divisome structure (Fig. 1B); we suspected that this region was inessential for PerM-FtsB binding and that it might destabilize Mtb FtsB in the absence of Mtb FtsL and/or FtsQ. In these constructs, expression was induced by anhydrotetracycline (ATc), which is solvachromatic, and we confirmed that there was no significant background signal in our imaging conditions (Supplementary Fig. S3). Deletion of FtsB^LQ^ clearly increased the level of membrane-localized FtsB, suggesting instability of Mtb FtsB in the absence of FtsL and FtsQ. However, FtsB instability in Mtb may result from other mechanisms [9], so we did not investigate this in depth. We did not observe any further effect on mTurquoise2 fluorescence when deleting FtsB^H^ in addition to FtsB^LQ^.

**Fig. 3.**
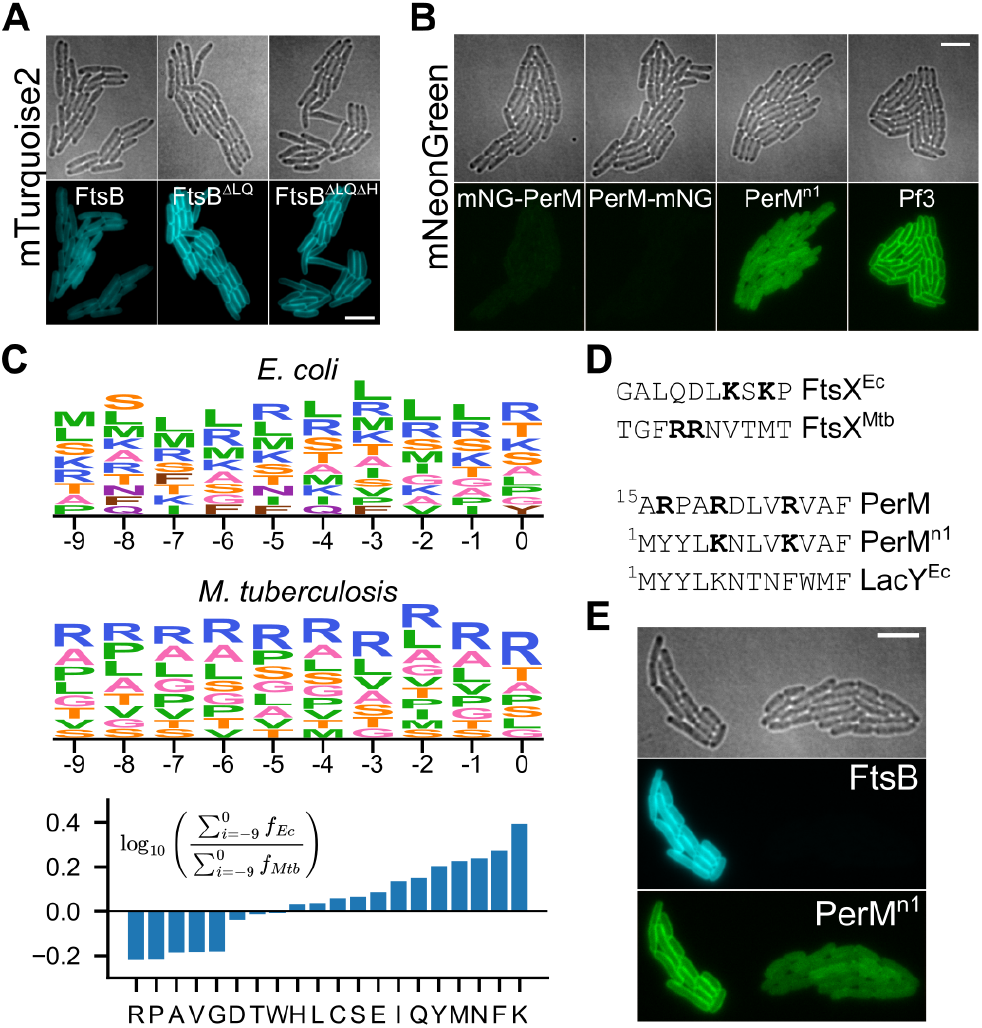
Expression of fluorescent fusion proteins with Mtb FtsB and PerM in *E. coli*. (**A**) Expression and membrane localization of FtsB and variants with deletions of FtsB^LQ^ or with deletion of both FtsB^LQ^ and FtsB^H^. Identical minimum and maximum intensities used to produce figures. (**B**) Expression and membrane localization for mNG-PerM, PerM-mNG, PerM^n1^-mNG, and Pf3-mNG. Identical minimum and maximum intensities Identical minimum and maximum intensities used to produce figures. (**C**) Top: residue frequency for positions preceding the first predicted transmembrane helix, inclusive of residues found at frequencies above 5%. Letter height is proportional to residue frequency with more frequent residues on top. Bottom: relative frequencies in *E. coli* and Mtb. (**D**) Top: residues preceding the first transmembrane helix in paralogs of FtsX, highlighting lysine and arginine residues. Bottom: comparison of PerM to PerM^n1^ and *E. coli* LacY. (**E**) Two adjacent microcolonies from a strain with co-expression of mTq2-FtsB and PerM^n1^-mNG. The microcolony on the right has coincidentally lost the plasmid encoding mTq2-FtsB, and exhibits reduced membrane localization of PerM^n1^-mNG. Scale bars 5 µm.

Conversely, initial attempts to construct mNeonGreen-labeled PerM failed for both N- and C-terminal fusion constructs (mNG-PerM and PerM-mNG; Fig. 3B). We manually inspected the predicted PerM structure as well as sequences of integral transmembrane protein orthologs in *E. coli* and Mtb, hypothesizing that an engineered construct, PerM^n1^, would exhibit higher expression. Specifically, we noticed an apparent “LK” motif in *E. coli* that rarely appeared in Mtb and was reminiscent of part of the twin-arginine translocation motif [26]. Our strategy in designing the PerM^n1^ construct was to modify PerM with aspects of the short LacY N-terminal sequence in ways that would not perturb the predicted PerM structure. While fluorescent protein expression was indeed much higher for PerM^n1^-mNG, it failed to exhibit the same degree of membrane localization as Pf3-mNG, which we used as a control given its efficient translocation and uniform transmembrane orientation [27].

We were surprised at how much the modifications in PerM^n1^ increased fluorescent protein expression levels and systematically investigated sequences in residues preceding the first predicted transmembrane helix in all proteins including at least one transmembrane helix in the *E. coli* and Mtb proteomes (Fig. 3C). We found large differences in the frequencies of some residues, most strikingly for differences in the relative frequencies of lysine and arginine. Fig. 3D shows how this is reflected in orthologous sequences of FtsX as well as in PerM^n1^ compared to Mtb PerM. Finally, we proceeded to co-express PerM^n1^-mNG and mTq2-FtsB and look for evidence of PerM-FtsB interaction. Serendipitously, one of two adjacent *E. coli* microcolonies in a co-expression experiment lost the plasmid for mTq2-FtsB protein expression, which occasionally happened without including selective antibiotics in agarose gel pads (Fig. 3E). We note that we used a YFP filter set for mNeonGreen imaging to avoid crosstalk from mTurquoise2. The large difference in PerM^n1^-mNG membrane localization between these two microcolonies suggested an unexpected role for FtsB in stabilizing PerM in the *E. coli* membrane.

### PerM stabilization in *E. coli* membrane by FtsB depends on FtsB^H^

In order to confirm that FtsB increases membrane-localized PerM protein expression and, if so, to investigate the role of FtsB^H^ in this phenomenon, we co-expressed mNeonGreen- and mTurquoise2-labeled constructs and analyzed fluorescence in *E. coli* microcolonies (Fig. 4). In these experiments, expression of PerM^n1^-mNG was induced with 100 µm IPTG. Localization controls mNeonGreen (strong cytoplasmic localization) and Pf3-mNG (strong membrane localization) were induced with 60 µm and 240 µm IPTG, respectively, to match typical PerM^n1^-mNG protein expression levels. Expression of mTurquoise2-labeled FtsB constructs was induced with 10nm ATc in all conditions except for one condition with no induction (indicated with down arrows in Fig. 4). Brightfield images of *E. coli* cells were segmented, and protein concentration was estimated to be proportional to total integrated fluorescence after subtracting background and normalizing by cell size.

**Fig. 4.**
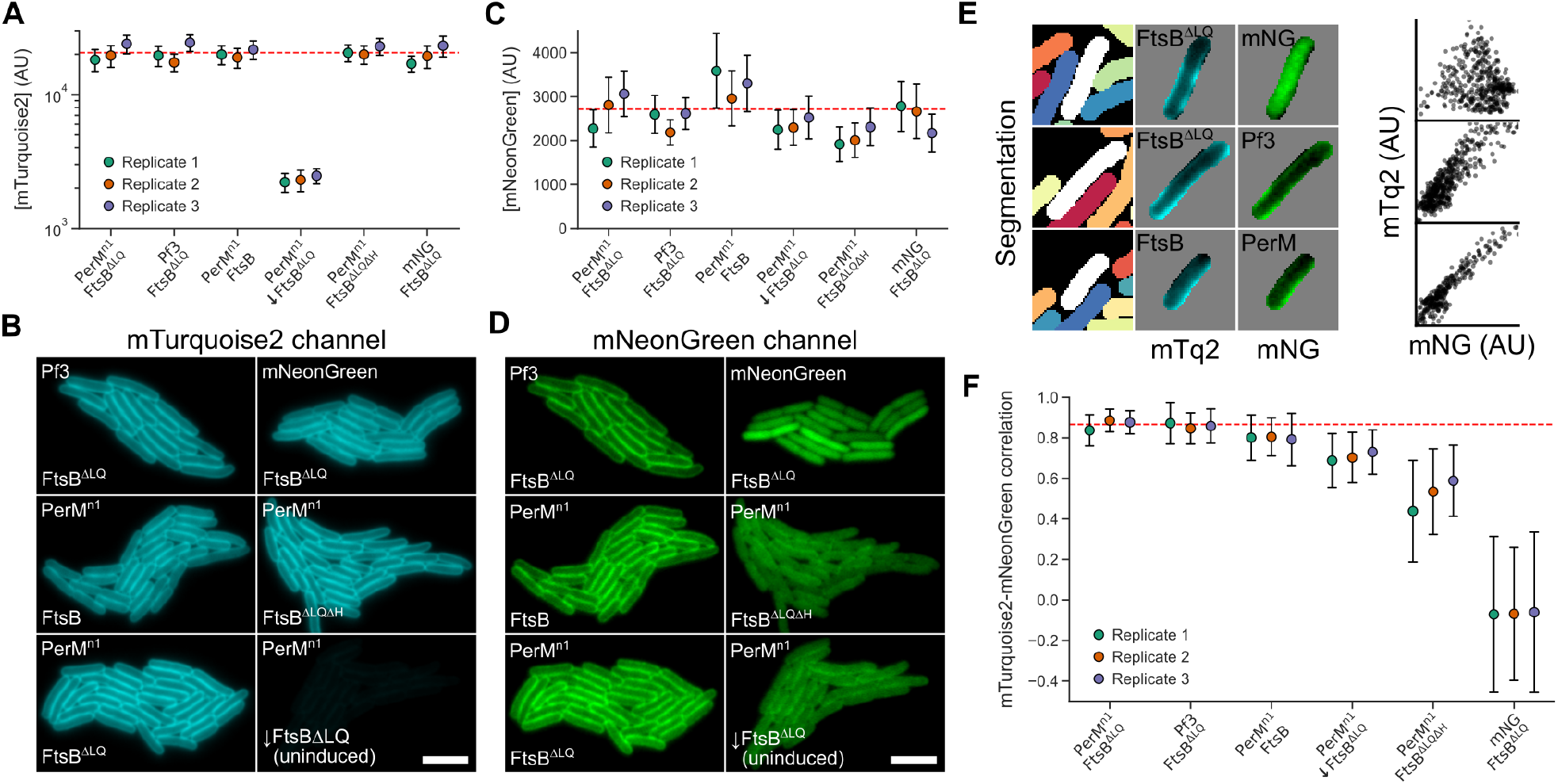
FtsB stabilizes PerM in the *E. coli* membrane. (**A**) Distribution of single-cell mTurquoise2 concentration (mean ± standard deviation) for FtsB constructs in different conditions, each with replicates. Dashed red line indicates the mean value for the reference condition (PerM^n1^-mNG, mTq2-FtsB^ΔLQ^, 100 µm IPTG, 10nm ATc). Uninduced FtsB^ΔLQ^ is indicated with a down arrow. A logarithmic scale is used to allow comparison of high- and low-protein-expression conditions. (**B**) Example data from microcolonies for each condition. The uninduced FtsB^ΔLQ^ condition has very low fluorescence in example data because identical minimum and maximum intensities were used. Scale bar 5 µm. (**C, D**) Distribution of single-cell mNeonGreen concentrations and example data for Pf3-mNG, mNeonGreen alone, or PerM^n1^-mNG in different conditions; prepared identically to **A** and **B**. (**E**) Left: example data analysis showing cell segmentation and isolated, single-cell intensities. Right: Scatter plots of single-pixel intensities for each cell show clear correlation for Pf3-mNG and PerM-mNG, but not for mNeonGreen alone. (**F**) Distribution of single-cell Spearman correlation coefficients (mean *±* standard deviation) for three replicates. Conditions ordered by decreasing mean correlation.

Relative to the reference condition (PerM^n1^ and FtsB^ΔLQ^, 10nm ATc), there was an 89% (*p* = 3.3 × 10^*−*12^) decrease in mTq2-FtsB^ΔLQ^ protein expression in the absence of induction, with no significant differences observed for other conditions. There was also no obvious difference in localization of any FtsB variant in any condition tested (Fig. 4B). The absence of any significant impact on full-length FtsB protein expression (*p* = 0.97) and the observation that PerM^n1^-mNG increases membrane localization of mTq2-FtsB (compare Fig. 4B and Fig. 3D to Fig. 3A) are consistent with the finding that PerM stabilizes FtsB [9] and that the relative level of PerM^n1^-mNG in this experiment is sufficient for FtsB stabilization. However, we did not explore this in detail given the lack of Mtb FtsL and FtsQ in our experimental system.

In contrast, there was a small reduction in PerM^n1^-mNG protein expression of 13.3% (*p* = 3.8 × 10^*−*2^) in the absence of induction of mTq2-FtsB^ΔLQ^ protein expression, and a reduction of 23.5% (*p* = 5.5 × 10^*−*4^) when FtsB^ΔLQ^ was replaced with FtsB^ΔLQΔH^. There was no significant change when FtsB^ΔLQ^ was replaced with wild-type FtsB (*p* = 0.29). This suggested that FtsB increased PerM stability in *E. coli*, and that this depended on the presence of FtsB^H^. However, inspection of Fig. 4C suggests that our sample size is overpowered (*N* = 10 930 cells in total from 3 replicates; 607 ± 44 cells per condition per replicate) given variation between replicates in mNeonGreen protein expression levels. Clearly, total fluorescence intensity did not capture the differences between conditions evident in Fig. 4D and an alternative method was required.

In preliminary experiments, we observed a large decrease in spatial correlation between mTurquoise2 and mNeonGreen fluorescence when FtsB^ΔLQ^ was depleted or replaced by FtsB^ΔLQΔH^ (Fig. 4D). The observation of small reductions in PerM^n1^-mNG levels despite apparently large impacts of different FtsB constructs on localization suggested that fluorescent mNeonGreen often remained in the cytoplasm following PerM^n1^-mNG degradation. We confirmed this by directly observing FtsB^H^-dependent degradation of PerM^n1^-mNG in cell extracts from samples differing in FtsB constructs and induction conditions; samples with mTq2-FtsB^ΔLQΔH^ and uninduced samples with mTq2-FtsB^ΔLQ^ had a greater fraction of a major low-molecular-weight band as well as a range of intermediate degradation products (Supplementary Fig. S4).

We hypothesized that quantifying spatial correlation would be a more sensitive measurement of PerM^n1^-mNG stability than mNeonGreen fluorescence, both less susceptible to variation in protein expression levels and robust against the presence of fluorescent, cytoplasmic PerM^n1^-mNG degradation products. Fig. 4E shows scatter plots of mNeonGreen and mTurquoise2 intensities for pixels in typical cells with cytoplasmic mNeonGreen, membrane-localized Pf3-mNG, and PerM^n1^-mNG. We calculated Spearman correlation coefficients from such distributions for segmented cells and Fig. 4F shows the mean and standard deviation of correlation coefficients for each condition and replicate.

In the reference condition (PerM^n1^-mNG, mTq2-FtsB^ΔLQ^, 10nM ATc), correlation was high (*ρ* = 0.87, 0.84–0.89 95% confidence interval) and indistinguishable from that when Pf3-mNG was expressed instead (*ρ* = 0.86, 0.82–0.90). All other conditions had significant reductions in correlation. Wild-type FtsB was marginally less well correlated (*ρ* = 0.80, 0.77–0.83). The apparent reductions in PerM^n1^-mNG protein expression levels discussed above were more strongly supported by analysis of spatial correlation, with a drop in correlation in the absence of mTq2-FtsB^ΔLQ^ induction (*ρ* = 0.71, 0.69–0.73) and a larger drop when mTq2-FtsB^ΔLQ^was replaced with mTq2-FtsB^ΔLQΔH^ (*ρ* = 0.52, 0.46–0.58). While FtsB^LQ^ was not absolutely essential for membrane localization of PerM^n1^, this was also observed in the absence of any FtsB expression (Fig. 3B).

### Single-molecule FtsB tracking reveals that FtsB^H^-dependent PerM binding

Our correlation analysis strongly suggested that FtsB interaction with PerM via FtsB^H^ is key for expression of PerM^n1^ protein in the membrane. Since we observed this for transgenic expression in *E. coli*, it is unlikely that PerM-FtsB interaction is mediated by host proteins. However, transient PerM-FtsB interaction could be sufficient for PerM stability without a large fraction of molecules being bound at any time. If PerM is a significant component in the core Mtb divisome as suggested by structure prediction, depletion phenotypes, and midcell localization [8, 9], we hypothesized that this implied long-lived PerM-FtsB interactions that could be detected by single-molecule tracking of FtsB diffusion. FtsB has a single transmembrane helix, while PerM is predicted to have 8 (Fig. 1A); based on the dependence of diffusion of integral membrane proteins on the number of transmembrane helices, we expected a reduction in the FtsB diffusion coefficient from approximately0.7 to 0.3 µm^2^ s^*−*1^ upon PerM binding [28].

We constructed mEos3.2-tagged variants of FtsB^ΔLQ^ and FtsB^ΔLQΔH^, and co-expressed them with either PerM^n1^-mNG or Pf3-mNG. Fig. 5A shows a typical frame from single-molecule imaging data, localized mEos3.2 molecules, and single-molecule tracks gathered in one movie showing that tracked molecules were detected near the middle plane of *E. coli* cells (approximately 0.5 µm from the microscope coverslip). In analysis of preliminary data, we found that analysis of either mean squared displacement or fitting jump-length distributions was sensitive to variable trajectory length. Thus, we utilized SASPT [29] to infer distributions of *apparent* diffusion coefficients using a method robust against short trajectories, variable localization error, and defocalization [30]. Note that we refer to an *apparent* diffusion coefficient, as diffusion within our images is largely confined to the one-dimensional perimeter at the middle plane and, further, confined at the cell poles. So, while this is sufficient to compare relative diffusion rates in different conditions, precise estimates of diffusion coefficients will require more sophisticated methods and would benefit from 3D tracking over the entire membrane.

**Fig. 5.**
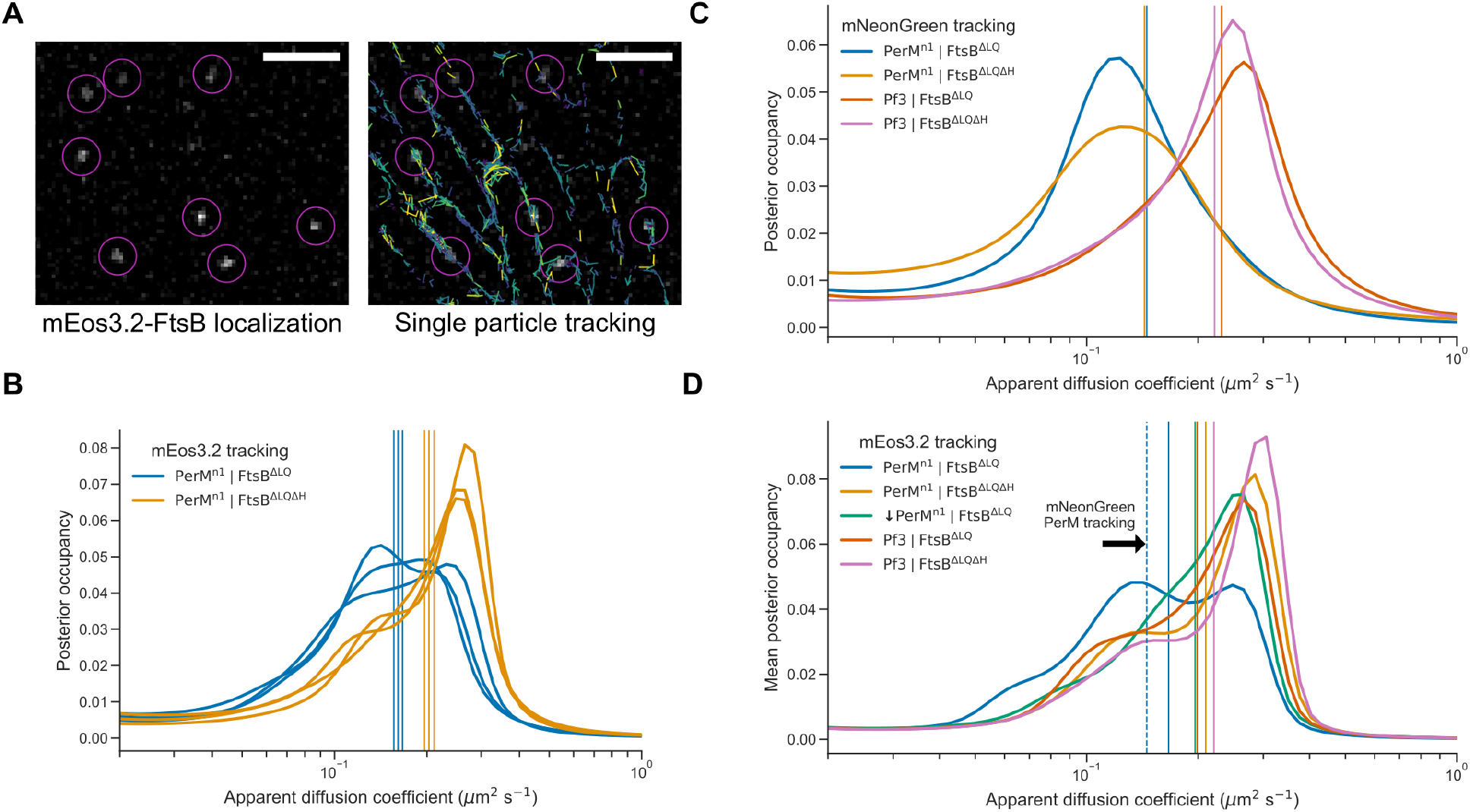
PerM-FtsB binding detected by FtsB single-molecule tracking. (**A**) Left: single, 33-ms frame from a fluorescence microscopy movie of mEos3.2-FtsB^ΔLQ^ diffusion in *E. coli*. Right: Single-molecule localization and tracking results for the entire movie show membrane-localized diffusion. Scale bar 2 µm. (**B**) Posterior occupancies in a model of regular Brownian motion and localization error were marginalized over localization error to estimate the distribution of apparent 2D diffusion coefficients. Distributions and their means (vertical lines) are shown for three replicates of experiments combining PerM^n1^-mNG with either mEos3.2-FtsB^ΔLQ^ or mEos3.2-FtsB^ΔLQΔH^. (**C**) Estimated distributions of diffusion coefficients inferred from single-molecule tracking of either PerM^n1^-mNG or Pf3-mNG with either FtsB^ΔLQ^ or FtsB^ΔLQΔH^ (single experiment). Vertical lines indicate the mean estimated diffusion coefficient for each condition. (**D**) Estimated distribution of diffusion coefficients for either mEos3.2-FtsB^ΔLQ^ or mEos3.2-FtsB^ΔLQΔH^ co-expressed with either Pf3-mNG or PerM^n1^-mNG (with or without addition of 100 µM IPTG), inferred from combining data from three replicates. Vertical lines indicate mean estimated diffusion coefficients and the dashed vertical line is replicated from **C**.

In tracking experiments, mEos3.2-FtsB constructs were expressed without induction so that PerM would be in excess and maximize potential binding. We inferred distributions of diffusion coefficients after marginalizing out localization error for three independent replicates (Fig. 5B). We observed reproducible differences in mean diffusion coefficients between FtsB^ΔLQ^ (0.161 ± 0.004 µm^2^ s^*−*1^; mean ± standard deviation) and FtsB^ΔLQΔH^ (0.203 ± 0.006 µm^2^ s^*−*1^; mean ± standard deviation). Distributions for both FtsB^ΔLQ^ and FtsB^ΔLQΔH^ suggested the presence of a slow-diffusion population at approximately 0.14 µm^2^ s^*−*1^ that is more prominent for FtsB^ΔLQ^. We also collected single-molecule tracking data for PerM^n1^-mNG and Pf3-mNG in single experiments. Despite limitations of this data (fewer localizations and variable spot density as mNeonGreen photobleaches over time), we inferred average diffusion coefficients of 0.144 µm^2^ s^*−*1^ and 0.226 µm^2^ s^*−*1^ for mNG-PerM and mNG-Pf3, respectively (Fig. 5C).

To estimate the most likely apparent diffusion coefficients, we inferred distributions from all data from each condition (Fig. 5D). This confirmed that mEos3.2-FtsB^ΔLQ^, when expressed with PerM, clearly exhibits slower diffusion than when PerM is replaced with Pf3 or in the absence of FtsB^H^. Peaks at approximately 0.14 and 0.25 µm^2^ s^*−*1^ are consistent with a substantial decrease in the rate of diffusion upon PerM binding. Compared to conditions with mEos3.2-FtsB^ΔLQΔH^ and conditions with Pf3, mEos3.2-FtsB^ΔLQ^ in the absence of induction of PerM^n1^-mNG exhibited the slowest diffusion (green line in Fig. 5D). This indicated that low-level PerM^n1^-mNG in the absence of induction has substantial mEos3.2-FtsB^ΔLQ^ binding. However, the effect size was clearly low and further work is required to determine the concentration dependence of this interaction.

### PerM-FtsB interaction could restrict conformational flexibility of the Mtb divisome

Our observation that FtsB^H^-dependent PerM-FtsB interaction was sufficiently long-lived to produce large changes in FtsB diffusion prompted us to explore how this interaction could impact the Mtb divisome. We conducted MD simulations to compare divisome constructs in which FtsB was truncated to FtsB^205^ (removing the uncertain C-terminal residues) or truncated to FtsB^185^ (additionally remove FtsB^H^). First, we performed MD for the Mtb divisome with PerM and full-length FtsB for 500 ns. FtsB N-terminal residues and other terminal residues of subunits without confident structure predictions were omitted. A period of initial MD was required since our structure prediction did not have all subunits at ideal relative orientations; this was especially significant for FtsQ since its transmembrane helix is distant from others. The final conformer after 500 ns was used to build systems with truncation to FtsB^205^ (FtsB^ΔC^) and to FtsB^185^ (FtsB^ΔCΔH^), and each system was simulated for 1 µs. The final conformer for FtsB^ΔC^ is shown in Fig. 6A.

**Fig. 6.**
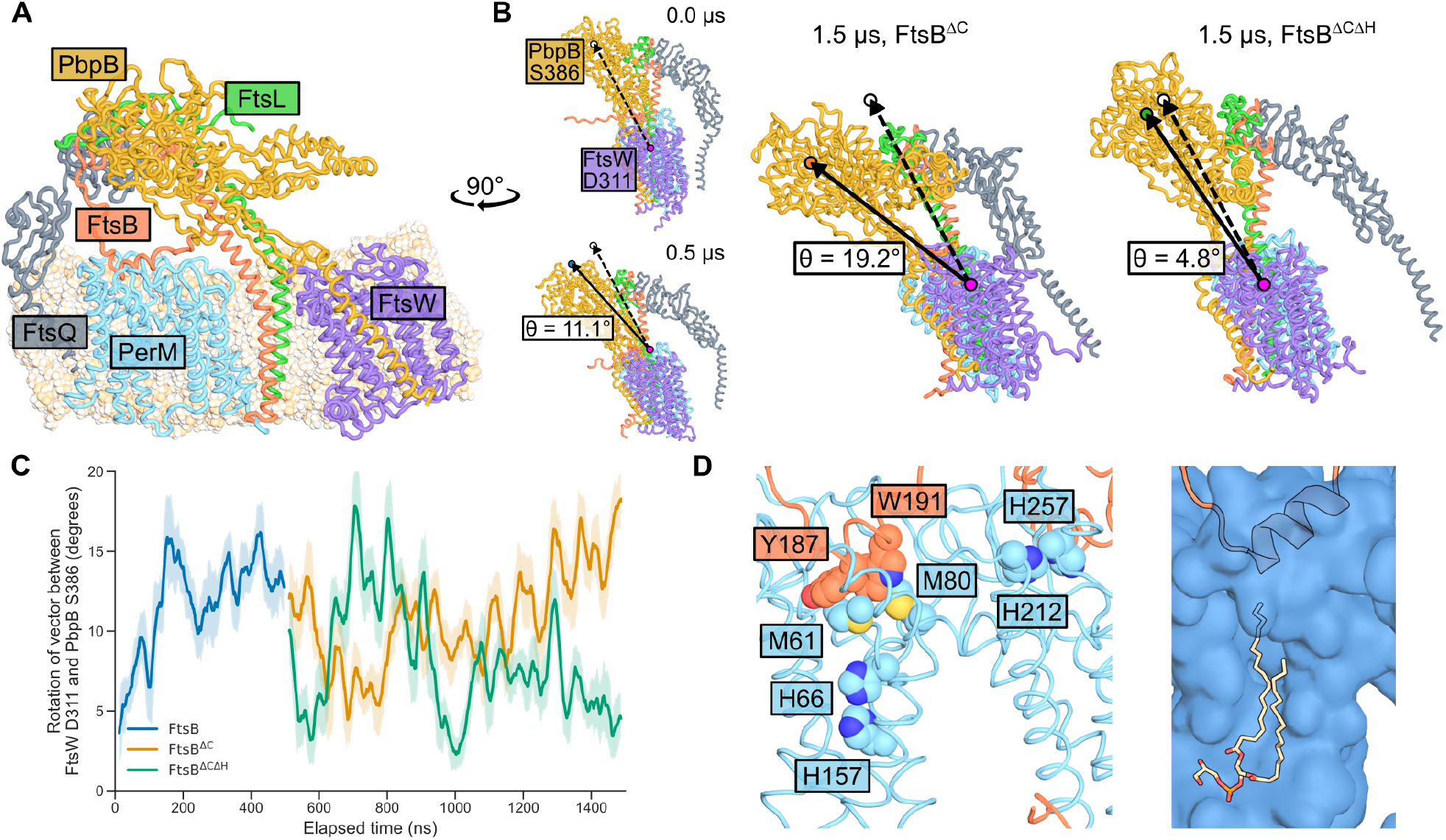
PerM-FtsB interaction constrains the Mtb divisome in MD simulations. (**A**) PerM simulated in context of Mtb divisome. Final conformer after 0.5 µs of MD with full-length FtsB followed by 1 µs of MD with FtsB^ΔC^. Note C-terminal truncation of FtsB relative to the predicted structure in Fig. 1A that is uncertain in this region. (**B**) Difference in tilt of PbpB transpeptidase domain relative to its conformation in structure prediction. Left: initial and final conformers of a 500 ns simulation including FtsB C-terminal residues. Locations of putative active-site residues FtsW^D311^ and PbpB^S386^ and the vector between them are highlighted; tilt of PbpB relative to FtsW is quantified by rotation of this vector. All conformers are aligned by FtsW C*α* atoms. The final conformer of the 500 ns simulation was used as the initial conformation for subsequent simulations. Right: Final conformers for FtsB^ΔC^ and FtsB^ΔCΔH^ simulations are shown along with their associated tilt angles. (**C**) Dynamics of tilt angles defined in **B** show that the FtsB^ΔCΔH^ simulation returned towards the elongated conformation observed in structure predictions. A 25 ns moving average of the tilt angle is shown and shaded areas indicate one standard deviation above and below the average angle. Line colors correspond to colors of circles in **B**. (**D**) Final conformer of PerM-FtsB aMD simulation after 1.1 µs. Left: residues in PerM near FtsB-binding interfaces suggest that PerM could potentially be sensitive to environmental cues. Right: a lipid often fills a channel in PerM in MD; here, in the final conformer of the aMD simulation, a phosphatidylglycerol molecule interacts with FtsB^H^.

Comparing the final conformers of the FtsB^ΔC^ and FtsB^ΔCΔH^ simulations, we examined the angle formed as the vector between the putative active site residues FtsW^D311^ and PbpB^S386^ rotates during MD simulation (Fig. 6B). This angle is a proxy for the tilt of the PbpB transpeptidase domain relative to FtsW that has been used to analyze Cryo-EM data as well as MD simulation. We observed a 19.2° tilt for FtsB^ΔC^, similar to that observed for *P. aeruginosa* FtsI comparing a structure prediction to its Cryo-EM structure [19] and for *E. coli* comparing a structure prediction to conformers following MD [20]. Conversely, with the loss of PerM-FtsB interaction mediated by FtsB^H^, the tilt angle reverted to a value within only 4.8° of the predicted structure. The progression of these angles over the course of MD is shown in Fig. 6C.

Relative to studies on the divisome in model organisms, there is a paucity of experimental data to relate our simulations to phenotypes associated with mutations to Mtb divisome components. Thus, we did not analyze interactions between Mtb divisome subunits in depth or conformations in and around FtsW and PbpB active sites. However, we note substantial differences the relative conformations of divisome components in final conformers for the FtsB^ΔC^ and FtsB^ΔCΔH^ simulations; tilt of the PbpB transpeptidase domain was associated with twisting of the PbpB head domain, FtsL, and FtsQ relative to FtsW. Lastly, in Fig. 6D we return to the final conformer of a PerM-FtsB simulation following aMD to highlight residues of interest in PerM that are suggestive of potential regulatory mechanisms. We also show how a phosphatidylglycerol molecule occupies a channel in PerM and interacts with FtsB^H^.

## Discussion

With our integrative approach combining fluorescence microscopy with molecular dynamics starting from predicted structures of protein complexes, we showed that FtsB^H^ directly mediates PerM-FtsB binding. Our results suggest that significance of PerM-FtsB interaction goes beyond FtsB stabilization [9] to potentially impacting the structure of the core Mtb divisome. Given the role of PerM in persistent Mtb infection and that it is essential in *M. smegmatis*, it will be interesting to learn whether FtsB also stabilizes PerM in mycobacteria, as it does in *E. coli*, and whether the PerM-FtsB interaction that we have identified can be targeted to disrupt regulation of cell division.

Our approach is straightforward to apply to other predicted protein-protein interactions. For example, Mtb PerM is located in the same operon as Rv0954 which similarly localizes to the Mtb divisome yet has no reported deletion phenotype [31]. Intriguingly, while PerM is associated with the divisome component FtsB, Rv0954 exhibited physical and genetic interactions with elongasome components such as RodA and PbpA. However, division and elongation machinery in mycobacteria are not perfectly analogous to those in other bacteria [32]. The cell-stress phenotypes of PerM depletion in Mtb suggest that conditional phenotypes may be identified for Rv0954 as well.

In Fig. 6D, we highlight PerM residues that could play roles in integrating environmental stress in proximity to FtsB^H^. These include buried histidines that are intriguing with respect to the Mg^2+^-dependent phenotype of PerM, suggesting sensitivity to nutrient depletion [8], methionine residues that make hydrophobic interactions with FtsB^H^ and could be sensitive to oxidation, and histidine residues near the membrane surface that could sensitize PerM-FtsB interaction to pH. Furthermore, in MD we routinely observed lipids to penetrate the first half of PerM and come into contact with FtsB, suggesting the possibility that bilayer composition can directly impact PerM-FtsB interaction.

In our experimental system, transgenic expression of Mtb PerM and FtsB proteins in *E. coli* makes it likely that we observed effects of direct PerM-FtsB interaction. This is an advantage of our system as it suggests that PerM-FtsB binding does not depend on phosphorylation or other post-translational modification specific to mycobacteria. While our single-molecule tracking experiments are not easily extensible to high-throughput investigation, PerM-FtsB and other Mtb protein-protein interactions can be explored by adopting our approach to develop FRET interaction reporters based on structures of predicted complexes. It will also be interesting to see whether our anecdotal success in engineering improved membrane protein expression in *E. coli* is applicable to other Mtb membrane proteins.

Our approach also carries the inevitable limitations of exploring an interaction in a surrogate system. It is critical to not extrapolate too much from our experiments before confirming results in Mtb. Our models of the Mtb divisome make countless predictions that are testable given the variety of tools for manipulating mycobacterial gene expression and recombineering. Additionally, our MD simulations omitted terminal residues in divisome components that are likely functional in some cases. This is only one example of interactions that we did not predict, but also cannot rule out. As another example, a specific FtsB-PerM interface in the membrane was not predicted and did not emerge in MD, but chimeras replacing FtsB™ with an alternative transmembrane helix could determine whether it contributes to PerM binding. One prediction arising from our work is that overexpression of FtsB^H^ in the Mtb or *M. smegmatis* periplasm could produce PerM-depletion phenotypes [9] via competitive inhibition of PerM-FtsB interaction.

Lastly, our observation that PerM-FtsB interaction constrains PbpB in a tilted conformation can be considered in light of results and discussion emerging from the Cryo-EM structure of the *P. aeruginosa* divisome [19], where an elongated conformation was considered to potentially reflect the catalytically active state. Recent work combining *in vitro* single-molecule and peptidoglycan polymerization assays with characterization of *in vivo* phenotypes supported this model for the gram-negative elongasome [33]. Within the context of this model and in light of our results, the role of PerM would be to reduce divisome activity by constraining PbpB in a conformation with relatively low transpeptidase activity. This is an apparent paradox given that PerM is essential in some conditions in Mtb and in *M. smegmatis*. However, the paradox can be resolved if PerM plays both a role in promoting FtsB stability (which can be bypassed with FtsB overexpression [9]) and also a conditionally essential role in regulating divisome activity that could contribute to persistent Mtb infection.

## Methods

### Structure prediction

PerM and FtsB sequences used for structure prediction are listed in Table 1 and additional Mtb sequences used for complex structure prediction are listed in Table 2. FtsB and PerM were identified in various actinomycete species and, after multiple sequence alignment, have between 19% and 61% (PerM) and 27% and 62% (FtsB) amino acid identity with Mtb orthologs. Protein complex structures were predicted using ColabFold [34]. Multiple sequences alignments were provided by MMseqs2 [35] using both paired and unpaired sequences in reference and environmental sequence databases. AlphaFold-Multimer [16] (Version 3, Model 4, 12 recycles) was used for structures compared in Fig. 1 and Supplementary Fig. S1 and for structure predictions utilized to build MD systems. PyMOL [36] was used for all structure visualizations and its `align` function was used to align FtsW structures and to calculate *RMSD* (using `cycles=0`).

**Table 1.**
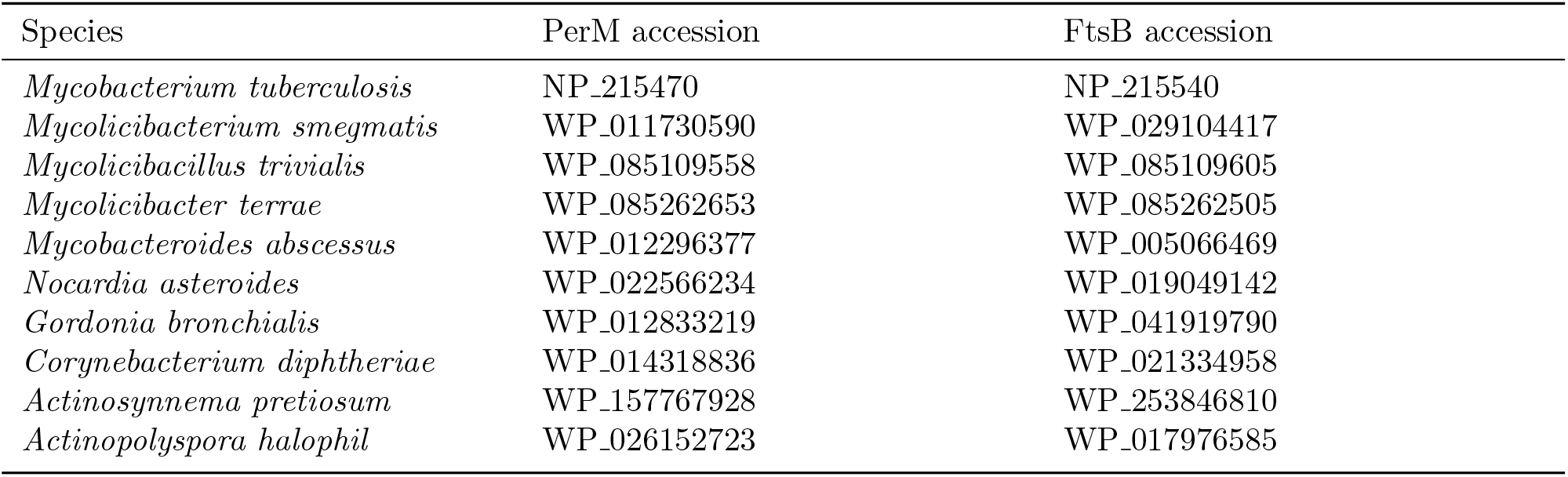
GenPept accession numbers for PerM and FtsB sequences used for different actinomycete species.

**Table 2.** GenPept accession numbers for PerM and FtsB sequences used for different actinomycete species.

| Protein | Accession |
| --- | --- |
| FtsQ | NP_216667 |
| FtsL | NP_216680 |
| FtsW | NP_216670 |
| PbpB | NP_216679 |

### Sequence analysis

To compare PerM and FtsB for different species, multiple sequence alignments were generated using Clustal Omega [37] and visualized using Biotite [38]. To compare amino acid frequencies at transmembrane helices, we used DeepTMHMM [39] to predict the positions of the first transmembrane helix for all proteins in reference proteomes for *E. coli* and Mtb (UniProt ID UP000000625 and UP000001584). Amino acid frequencies and corresponding sequence logos for the 10 residues preceding the first residue in the first predicted transmembrane helix were calculated and plotted using Logomaker [40].

### Molecular dynamics

Molecular dynamics systems were constructed from structures of predicted complexes (or from the final conformer of the initial 500 ns divisome complex simulation) with the CHARMM-GUI Membrane Builder [41]. Shared simulation parameters chosen in CHARMM-GUI were: 150 mm KCl, CHARMM36m forcefield [42], 310 K temperature, NPT ensemble, and capping all non-native N- and C-terminal residues with ACE and CT3, respectively. Scripts for OpenMM [43] output by CHARMM-GUI were modified and OpenMM was used for MD simulation. Simulation system sizes are detailed in Table 3.

**Table 3.** Number of total atoms and numbers and types of lipid residues in MD systems.

| System | Atoms | Residues |  |
| --- | --- | --- | --- |
|  |  | POPE | POPG |
| PerM-FtsB | 156168 | 210 | 70 |
| PerM only | 105155 | 213 | 71 |
| Complex (FtsB) | 724208 | 1246 | — |
| Complex (FtsB<sup>ΔC</sup>) | 709754 | 1239 | — |
| Complex (FtsB<sup>ΔCΔH</sup>) | 709645 | 1239 | — |

Simulation systems differed in a few ways (other than small differences in residues included, shown in Table 4). First, bilayers simulations for the PerM-FtsB complex and PerM alone included a 3:1 POPE:POPG ratio commonly used for simulations of *E. coli* membranes since MD results were compared to *E. coli* microscopy data. Simulations for divisome complexes used pure POPE bilayers, and future work may investigate differences in more complex model membranes [44]. Second, simulations of the PerM-FtsB complex and PerM alone followed a three-stage aMD protocol described in the main text with 2 fs timesteps. For the aMD stage, OpenMM scripts from CHARMM-GUI [45] were modified to utilize empirically determined dual boost potentials as previously described for MD of transmembrane proteins [46] and to output a log of applied boost potentials. We monitored boost potentials applied to dihedral angles and total energy and found the magnitudes of the two boost potentials were similar. For the final two simulations depicted in Supplementary Fig. S2, the *λ* parameter used to calculate the dihedral boost reference energy and acceleration factor was reduced from 0.3 to 0.15. Lastly, divisome complex simulations utilized hydrogen mass repartitioning [47] and 4 fs timesteps.

**Table 4.**
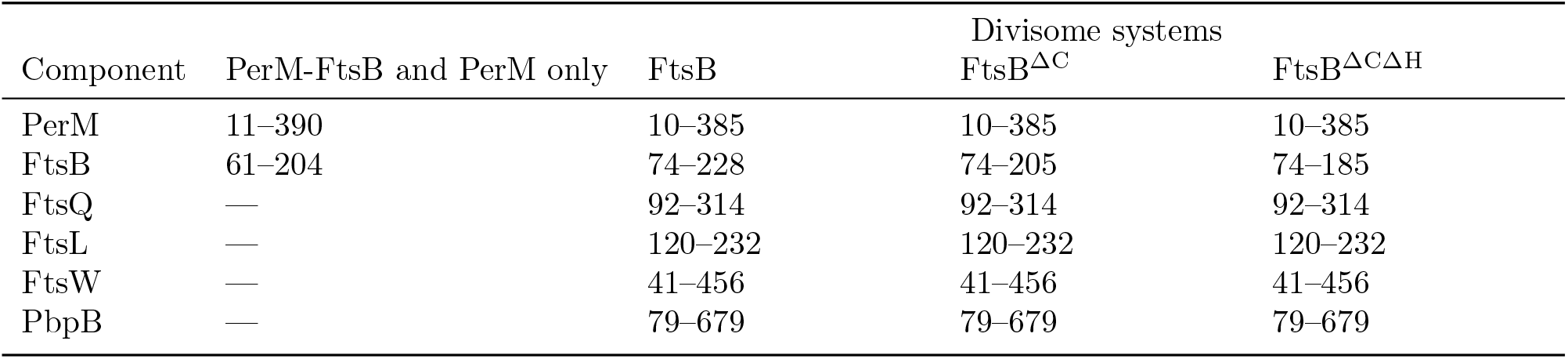
Residues included in MD simulation systems.

Atoms of interest for analysis shown in Fig. 2B and Fig. 6C were identified by investigating sequence alignments to orthologs of FtsB, PbpB, and FtsW to determine conserved FtsB^H^ residues and putative activesite residues. The MDAnalysis package [48, 49] was used to extract trajectories wrapped around the center of mass of protein atoms, to align divisome complex trajectories to FtsW^60–407^ (omitting more mobile cytoplasmic residues), and to extract atomic coordinates. All-atom coordinates of initial and final conformers as well as protein-only trajectories are available as supplementary data.

### *E. coli* strain construction

In addition to Mtb PerM and FtsB (and variations described in the main text), capsid protein G8P from bacteriophage Pf3 (UniProt identifier P03623; referred to as “Pf3” elsewhere in this manuscript) was utilized as a control based upon its small size, efficient translocation, and uniform transmembrane orientation [27]. Proteins were expressed as translational fusions with fluorescent proteins mTurquoise2 [25], mNeonGreen [24], or mEos3.2 [50]. No linkers were added between Mtb protein sequences and fluorescent proteins as we expected that native termini were disordered based on structure predictions. Sequences for all proteins were codon-optimized for *E. coli* protein expression. Plasmids for FtsB-expressing constructs were constructed by isothermal assembly from low-noise, tetracycline-inducible, ampicillin-resistant vectors [51]. Plasmids for expression of mNeonGreen as well as mNeonGreen fusions to PerM variants and Pf3 were constructed from IPTG-inducible, spectinomycin-resistant vectors [52]. All plasmids and induction conditions used in this study are summarized in Table 5.

**Table 5.** *E. coli* plasmids utilized in this study and their respective induction conditions. Mutations are described relative to wild-type Mtb protein sequences in Table 1 and Table 2.

| Plasmid | Selection | Induction | Expressed protein | Mutation |
| --- | --- | --- | --- | --- |
| pJRF002 | Carbenicillin | 10 nM ATc | mTq2-FtsB | — |
| pJRF007 | Carbenicillin | 10 nM ATc | mTq2-FtsB<sup>ΔLQ</sup> | FtsB<sup>Δ111–163</sup> |
| pJRF007<sub>helix</sub> | Carbenicillin | 10 nM ATc | mTq2-FtsB<sup>ΔLQΔH</sup> | FtsB<sup>Δ111–163Δ186–194</sup> |
| pZH904 | Carbenicillin | 10 nM ATc | mEos3.2-FtsB<sup>ΔLQ</sup> | FtsB<sup>Δ111–163</sup> |
| pLG906 | Carbenicillin | 10 nM ATc | mEos3.2-FtsB<sup>ΔLQΔH</sup> | FtsB<sup>Δ111–163Δ186–194</sup> |
| pJRF001 | Spectinomycin | 100 μM IPTG | mNG-PerM | — |
| pJRF004 | Spectinomycin | 100 μM IPTG | PerM-mNG | — |
| pJRF004<sub>n1.cWT</sub> | Spectinomycin | 100 μM IPTG | PerM<sup>n1</sup>-mNG | PerM<sup>Δ1–23</sup>:MYYLKNLVK |
| pJRF010 | Spectinomycin | 240 μM IPTG | Pf3-mNG | — |
| pZH813 | Spectinomycin | 60 μM IPTG | mNeonGreen | — |

Plasmids were transformed into *E. coli* strain MG1655 and selected for growth on 50 mg L^*−*1^ carbenicillin and/or spectinomcin in LB or SOB media. For microscopy experiments, cells were grown overnight from LB plates in rich defined medium [53] with 0.2% glucose (ForMedium Neidhardt Basal Salt Mixture, Glucose 20 g L^*−*1^, and Neidhardt Supplement Mixture, Complete), adjusted to pH 7.0 with KOH. All cell growth and imaging took place at 37 ° C. Consistent with our earlier work, our plasmids allowed for independent induction of co-expressed genes [52]. In preliminary experiments, we confirmed that mTurquoise2- and mNeonGreen-labeled proteins could be independently induced, that inducing protein expression did not impact growth at expression levels utilized in this work, and that there was no significant crosstalk between mTurquoise2 (CFP filter set) and mNeonGreen (YFP filter set).

### Fluorescence microscopy

All imaging was done on a Leica DMI6000 inverted microscope using illumination from either a Leica EL6000 source or with laser excitation at 514nm (mNeonGreen tracking) or combined 405nm and 561nm laser excitation (mEos3.2 tracking), adjusting 405nm laser intensity to achieve acceptable densities of photoactivated mEos3.2 molecules. Data was acquired using a 100×/1.46 a-plan apochromat oil immersion objective with additional 1.6× magnification, Leica type F immersion oil, and an Evolve 512 EM-CCD camera (Photometrics), giving a 100nm pixel size and a narrow depth of field. Fluorescence images were acquired with three different filter sets: Chroma 49001 (mTurquoise2), Semrock LF514-B (mNeonGreen), and a Chroma custom dual-laser filter set (mEos3.2; ZET405/561x, ET610/75m, ZT405/561/657rpc-UF2). Images of example data were composed using Fiji [54], with linear scaling and maintenance of minimum and maximum intensity values for all comparable images.

Microscope samples were prepared by sandwiching agarose gel pads (rich, defined media with 2% Invitrogen UltraPure agarose) of approximately 100 µL between microscope slides and acid-cleaned #1.5H coverslips (Marienfeld). For all experiments, samples were reinoculated from overnight cultures at a 1:1000 dilution in media including IPTG and/or ATc at specified concentrations, grown for 3 h, added to a gel pad prepared from the same media, and grown for an additional 1.5 h before acquiring data. Microscope sample temperature was held at 37 ° C using a combination of a heated microscope sample chamber and an objective heater. This protocol was based on empirical observations that cells reached steady state fluorescence and growth in less than 1 h after sample preparation, and that excessive cell density led to reduced growth and fluorescence after approximately 3 h. In preliminary experiments, we also noted that this combination of growth media and agarose exhibited high background with substantial variation between experiments, found that this could be attributed to agitation while melting agarose. We subsequently minimized agitation while preparing gel pads.

To obtain comparable data for constructs expected to have different translation initiation rates, we conducted preliminary experiments and determined that 60 µm and 240 µm IPTG induction for plasmids expressing mNeonGreen and Pf3-mNG, respectively, were approximately equivalent to 100 µm IPTG for plasmids expressing PerM^n1^-mNG. Protein expression was always induced with IPTG and ATc concentrations listed in Table 5 except in the uninduced conditions indicated by down arrows in figures. To increase reproducibility in fluorescence correlation and single-molecule tracking analysis, we aimed to image the middle plane of *E. coli* cells approximately 500 µm from the microscope coverslip. For fluorescence correlation analysis, we obtained images of 20 fields of cells for each condition in each of 3 independent replicates. For single-molecule tracking analysis, we obtained streaming movies (33 ms exposure time) of 5 fields of cells for each condition in each of 3 independent replicates (except for mNeonGreen tracking for which we did not replicate the experiment). Replicates were obtained on separate days, and sample preparation steps were staggered in time to minimize differences in how samples were handled for each condition.

### Fluorescence correlation analysis

Brightfield images were segmented using Omnipose [55]. The low and variable contrast in our brightfield data produced variation in widths of segmented objects, so objects were morphologically thinned to single-pixel width and then thickened to a uniform width that was more consistent with observed cell morphology. Subpixel channel alignment was achieved by maximizing pairwise cross-correlation of low-pass filtered fluorescence images with inverted brightfield images (*i*.*e*. mTurquoise2 and mNeonGreen images were aligned to brightfield images—not to each other). Background intensity was estimated as the mode of a histogram of low-intensity pixels outside of *E. coli* cells; in all conditions including uninduced mTq2-FtsB^ΔLQ^, cellular autofluorescence was very low relative to signal from fluorescent proteins. Fluorescence intensities in pixels corresponding to segmented cells after alignment and background subtracted were compared as described in the main text.

For mNeonGreen and mTurquoise2 concentration and correlation comparisons, a mixed linear model was fit to data points from single cells with random slopes and intercepts to allow for day-to-day variation. Two-tailed p-values were calculated for comparison to the reference condition (PerM^n1^ and FtsB^ΔLQ^, 100nm ATc) and adjusted for multiple comparisons using the Holm-Šidák method. Effect sizes are reported for significant results only (adjusted *p* < 0.05). Unless stated otherwise, errors in the manuscript are 1 standard error of the mean. We followed our designed analysis protocol and manually screened data to identify and exclude image fields that could contribute to outliers (large shifts in sample position, poor focus, or unusual levels of fluorescent protein expression for one or both proteins in a microcolony; 32 of 360 total images) and also excluded cells falling outside the middle 95th percentile in either fluorescence intensity or cell area. However, we found that removing all data filtering steps did not significantly impact differences we observed between experimental conditions.

### Direct observation of FtsB-dependent PerM^n1^-mNeonGreen degradation

Strains expressing PerM^n1^-mNeonGreen together with either mTurquoise2-FtsB^ΔLQ^ or mTurquoise2-FtsB^ΔLQΔH^ were cultured overnight in 1mL of rich defined medium supplemented with 50 mg L^*−*1^ spectinomycin and carbenicillin. Overnight cultures were diluted 1:1000 into 5mL of fresh media of the same composition with the addition of different inducers (100 µm IPTG, 10nm ATc, or both). After a 3 h incubation at 37 ° C, the cells were harvested by centrifugation (8000 g, 1 min, 4 ° C) and washed twice with pre-cooled PBS. Following the final wash, the cells were resuspended in lysis buffer (PBS, 1 mg mL^*−*1^ lysozyme, 1×cOmplete protease inhibitor cocktail (Roche), 1% Triton X-100) and sonicated for 10 min using in cycles of 30 s on and 30 s off using a Bioruptor Plus sonication device. Cell lysates were mixed with 3×sample loading buffer (187.5 mm Tris, 6% SDS, 30% glycerol, 0.03% bromophenol blue, pH 7) and 30×DTT (1.25m). Samples were not heated in order to maintain mNeonGreen fluorescence [56]. Proteins were separated by Tris-Glycine-SDS electrophoresis (12% acrylamide) together with a prestained ladder (Bio-Rad Precision Plus Dual Color). The gel was rinsed in water and imaged using an iBright FL1500 to observe mNeonGreen and mTurquoise2 signals in two channels. One channel detected both mTurquoise2 and mNeonGreen (455nm to 485nm excitation and 508nm to 557nm emission) and the other channel detected mNeonGreen with extremely low mTurquoise2 efficiency (490nm to 520nm excitation and 568nm to 617nm emission). The relative degree of PerM^n1^-mNeonGreen degradation was estimated as the fraction of the major high- and low-molecular weight bands observed in the low-molecular weight band compared to the mTurquoise2-FtsB^ΔLQ^ condition with 10nm ATc.

### Inference of diffusion coefficient distributions

We extracted single-molecule trajectories from image stacks using TrackMate [57]. Spots were detected using `DogDetector` (250nm spot radius) and tracked using `SimpleSpareseLAPTracker` (500nm maximum linking distance). Thresholds were determined empirically for mEos3.2 (160) and mNeonGreen (200) data, and the first 250 frames of mNeonGreen movies were omitted to exclude data before sufficient photobleaching had occurred to acquire single-molecule tracks of mNeonGreen molecules. Trajectories were 3.56 ± 0.03 and 3.65 ± 0.11 frames long for mEos3.2 and mNeonGreen, respectively (mean and standard error of the means for 15 mEos3.2 and 4 mNeonGreen experiments; 5 streaming movies acquired in each experiment). We then used SASPT [29] to infer posterior distributions of track diffusion coefficients and localization errors, marginalizing out localization distributions for analysis related to Fig. 5. Posterior distributions were inferred for log-spaced diffusion coefficients between 0.01 and 10 µm^2^ s^*−*1^. Distributions were normalized to occupancies between 0.02 and 1 µm^2^ s^*−*1^ to exclude states with highly variable occupancy at the extremes of the diffusion coefficient range.

Our primary analysis used default SASPT parameters (200 iterations of refining state occupancy, trajectories longer than 10 frames split into trajectories 10 frames long or less) and we increased the `sample_size` parameter to avoid subsampling data. Based on our observation that single-molecule tracks clearly show the perimeter of *E. coli* cells at the middle plane, we used 500nm for the `focal_depth` parameter. We also note that we manually inspected maximum projections of all image stacks to verify that cells were well focused. To check for sensitivity to SASPT analysis parameters, we repeated posterior occupancy inference with various conditions: (1) excluding trajectories that were relatively long (> 10 frames which can arise from fluorescent objects that do not photobleach) or short (2-frame trajectories that can arise from false positive localizations), (2) fixing localization error to 25nm, and (3) increasing state array refinement to 1000 iterations. We found that SASPT results were robust against parameter choices and that the largest difference was somewhat narrower distribution peaks with increased iterations.

## Acknowledgements

ZH is supported by Fundação para a Ciência e a Tecnologia (FCT) through MOSTMICRO-ITQB (DOI 10.54499/UIDB/04612/2020; DOI 10.54499/UIDP/04612/2020) and LS4FUTURE Associated Laboratory (DOI 10.54499/LA/P/0087/2020). ZH and RX are supported by a MOSTMICRO-ITQB Exploratory Projects grant. ZH and JRF thank Sara F. Costa for assistance and insight on this project. ZH thanks Lúısa Guerreiro and Margarida Pereira who worked on this project during internships in our lab.

## Data availability

Plasmids described in this manuscript will be made available through AddGene. Data and analysis notebooks are available on request and will be made available upon publication at https://github.com/smmlab/mtb-perm-ftsb. Raw microscopy data and MD trajectories (protein atoms only) will be deposited at the Zenodo repository shortly and can be found by visiting the URL above.

## Author contributions

JF: molecular cloning, data acquisition, preliminary data analysis and interpretation. RX: data acquisition, methods development, data analysis. ZH: structure predictions and simulation, data acquisition, data analysis. All authors wrote and edited the manuscript.

## Supplementary Figures

**Fig S1.**
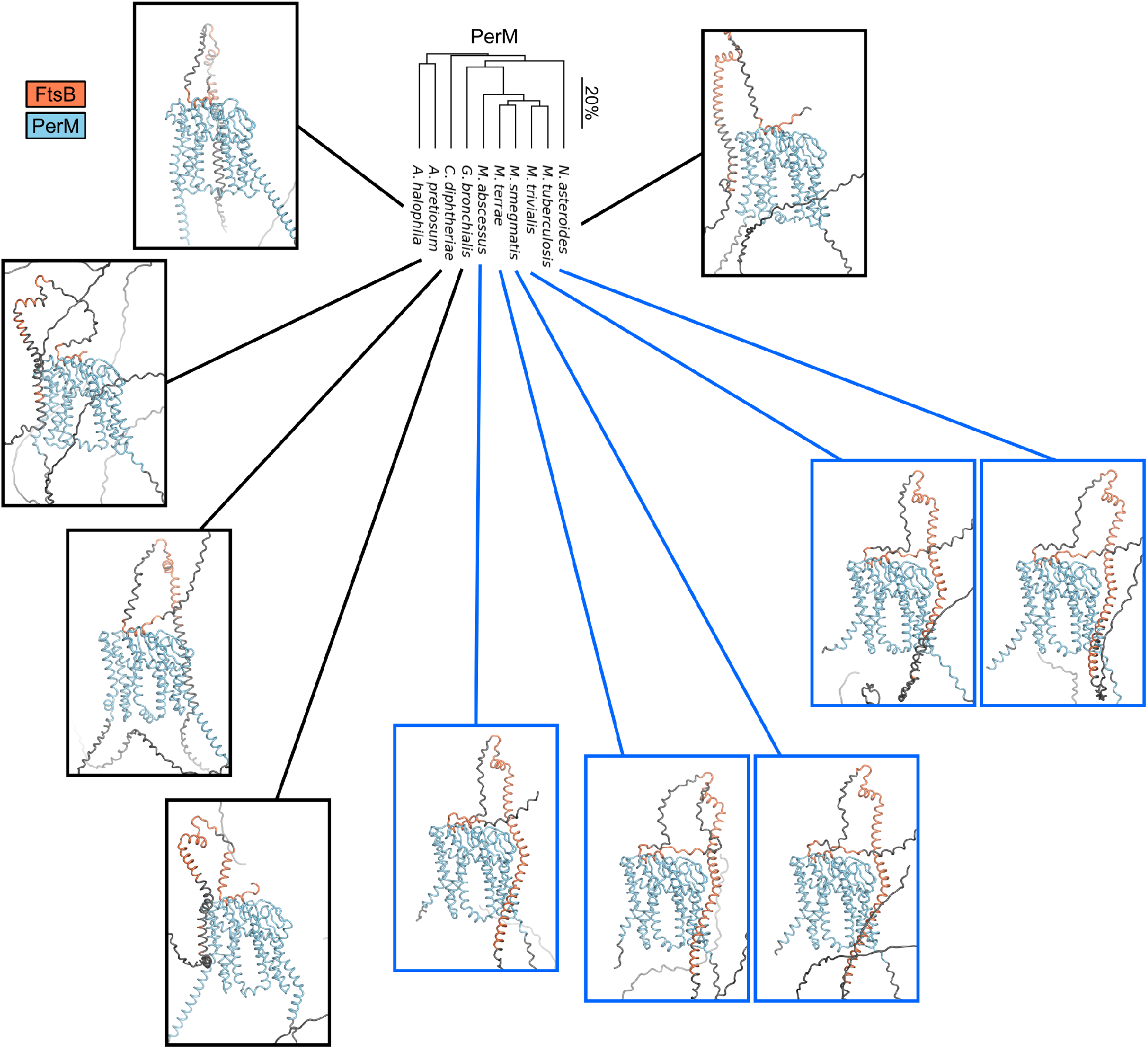
The PerM phylogenetic tree from Fig. 1 is reproduced and compared to PerM-FtsB complexes predicted from full-length sequences for the same species. The predicted orientation of FtsB™ relative to PerM observed for Mtb is only observed for more closely related species (blue lines). However, FtsB^H^ interaction with PerM is predicted for all species. Residues with *pLDDT* < 50 are colored gray.

**Fig S2.**
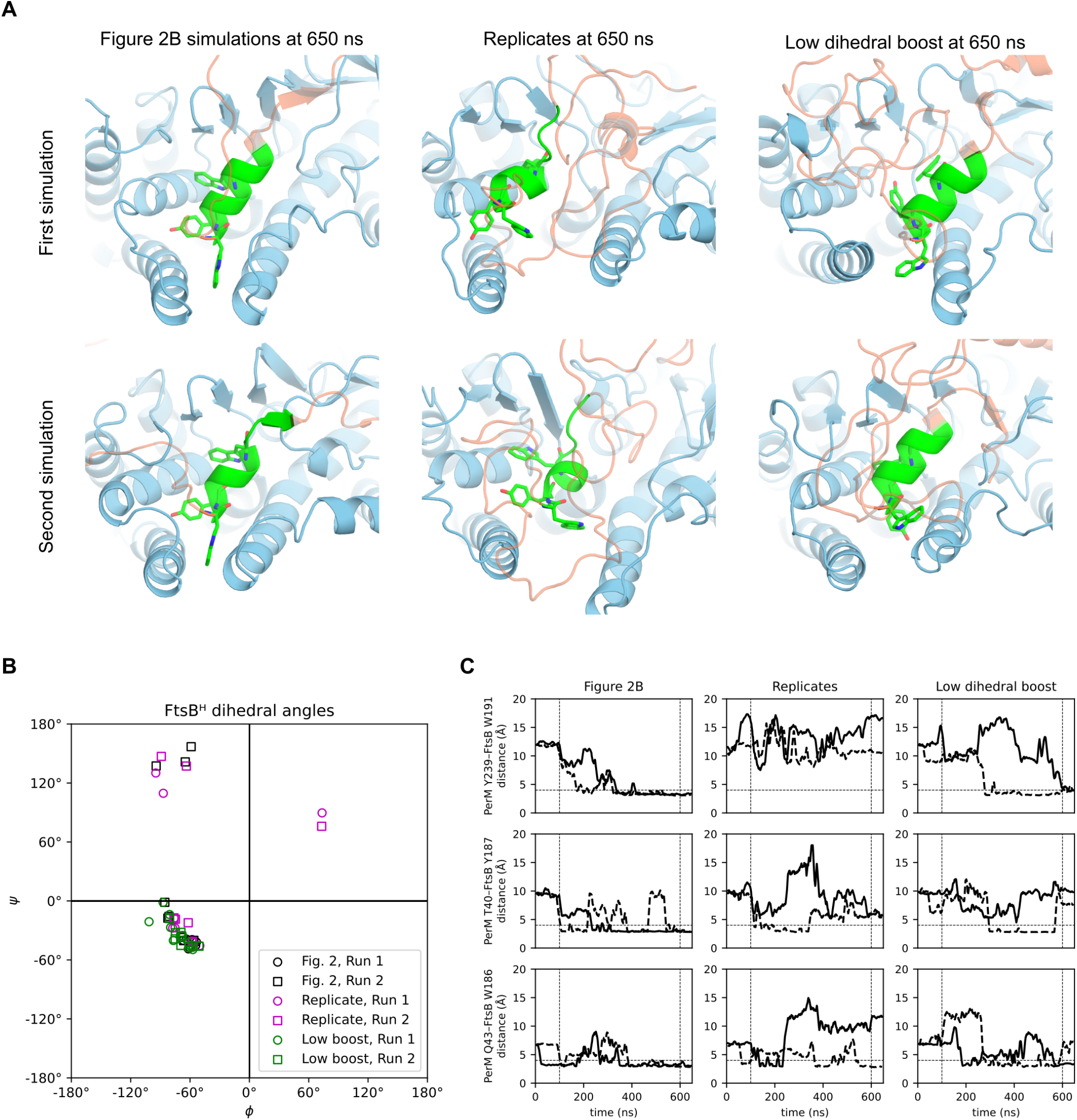
(**A**) Left: conformers at 650 ns for the PerM-FtsB aMD simulations described in the main text; FtsB^H^ residues are shown in green with a licorice representation for residues W186, Y187, and W191. Other FtsB residues are transparent orange and PerM is blue. FtsB^H^ secondary structure is disrupted in the second simulation. Middle: final conformers in two additional 650 ns replicate simulations. FtsB^H^ secondary structure is disrupted in both simulations. Right: final conformers in two 650 ns simulations in which *λ* value used to define dihedral boost parameters is reduced from 0.3 to 0.15. FtsB^H^ secondary structure is maintained in both simulations. (**B**) Ramachandran plot for FtsB^H^ residues in 650 ns conformers shows that *α*-helical dihedral angles are maintained for the first simulation in Fig. 2 and for both low boost simulations, but lost for residues in other simulations. (**C**) Left: reproduction of Fig. 2B plotted through 650 ns; a horizontal line is added at 4 °A. Middle: In replicate simulations that lose FtsB^H^ *α*-helical secondary structure, hydrogen bonds identified at the PerM-FtsB interface rarely form and a bond never forms between PerM Y239 and FtsB W191. Right: These hydrogen bonds form more frequently when the dihedral boost is reduced.

**Fig S3.**
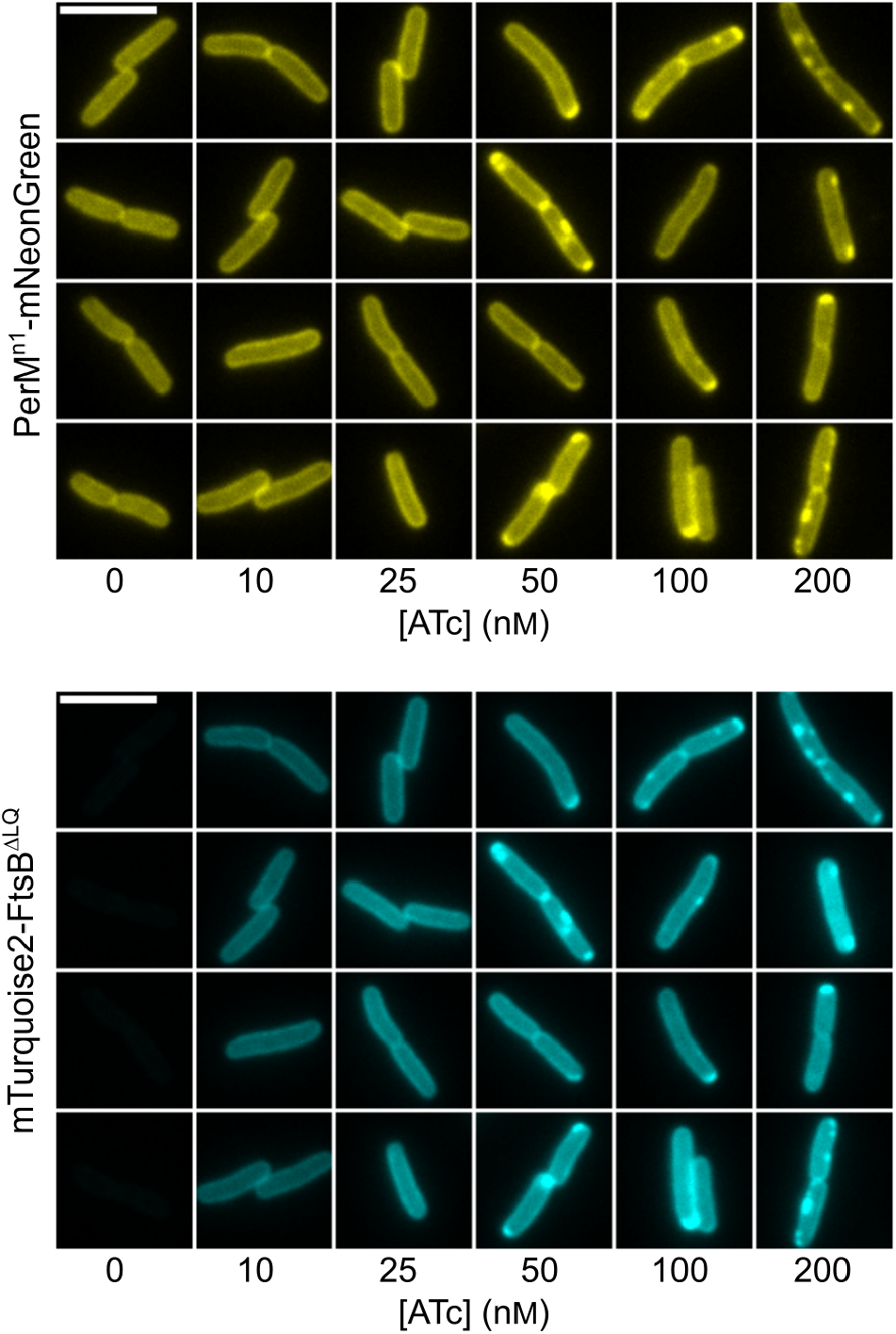
ATc does not add substantial background to either mTurquoise2 or mNeonGreen images. Cells with plasmids expressing PerM^n1^-mNeonGreen and mTurquoise2-FtsB^ΔLQ^ were grown in the same conditions used for experiments reported in the main text with 100 µm IPTG and with ATc at concentrations up to 20×the maximum concentration used in analyzed data. All images for each channel are shown with the same minimum and maximum intensity scaling to facilitate comparison of background and cellular fluorescence intensity. ATc concentrations at and above 50nm (5×the concentration used in our experiments) led to aggregation and a lack of further mTurquoise2 fluorescence increase outside of aggregates; aggregation of PerM^n1^-mNeonGreen as well suggests that membrane protein translocation is beyond capacity for this strain and growth condition. No increase in background was observed in either channel at high ATc concentrations. Four regions of interest are shown for each condition with the same regions shown for both mNeonGreen and mTurquoise2 images.

**Fig S4.**
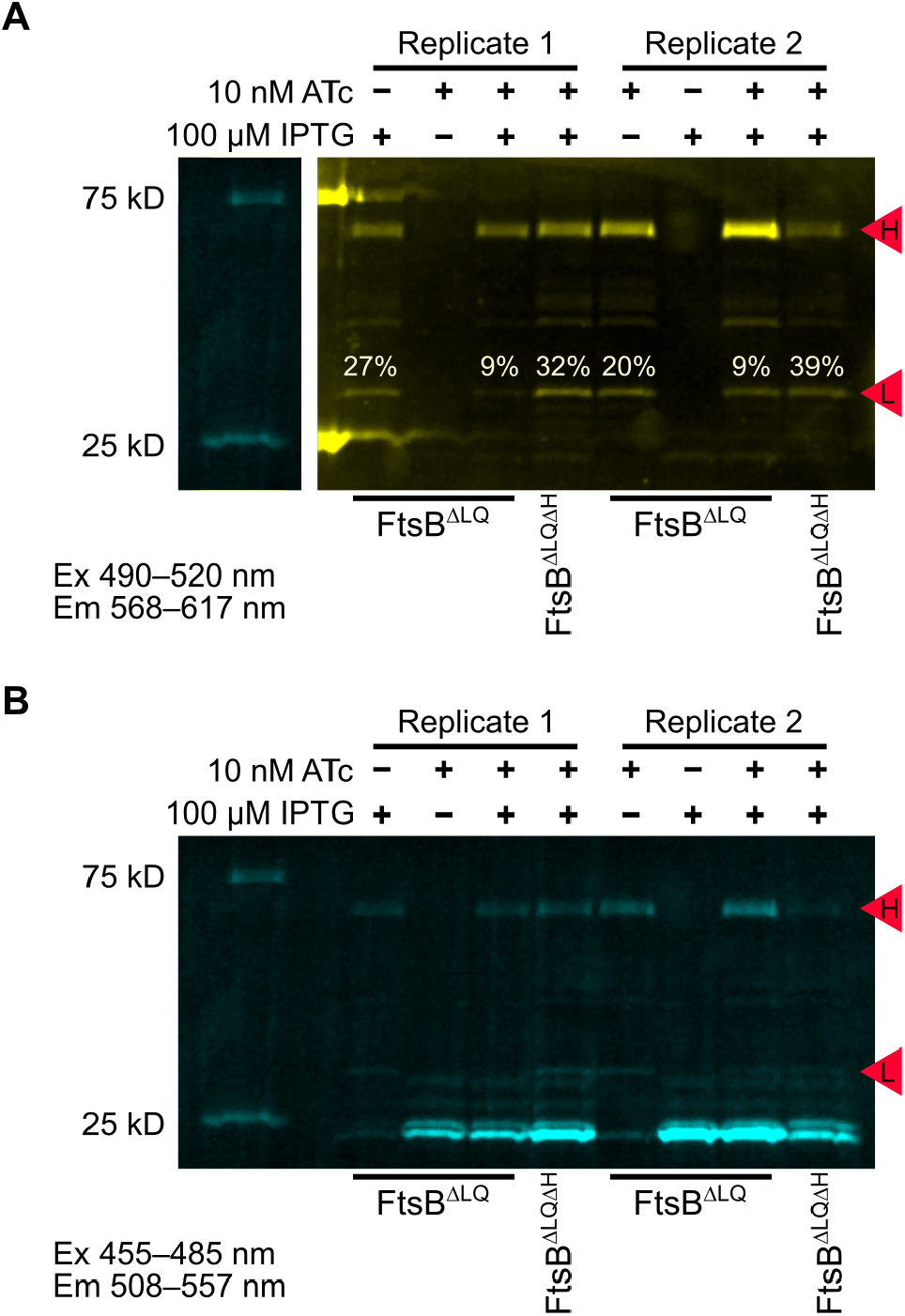
Direct observation of PerM^n1^-mNeonGreen degradation by fluorescent SDS-PAGE. (**A**) Cell extracts for strains that all include a plasmid expressing PerM^n1^-mNG as well as mTq2-FtsB constructs with deletions corresponding to labels for each lane were grown in different induction conditions and separated by SDS-PAGE in conditions that maintain mNeonGreen and mTurquoise2 fluorescence. Degradation of PerM^n1^-mNG was quantified in two replicates of cell extracts as the integrated band intensity of the major degradation product (labeled L) relative to the sum of intensities for band L and band H, which has a molecular weight approximately corresponding to that of PerM^n1^-mNG (71 kD). Intermediate degradation products were not quantified. A weak band with lower molecular weight also appears in the absence of IPTG, and corresponds to a protein with mTurquoise2 fluorescence that is inefficiently detected in this channel. The ladder band is replaced with that from the blue/green image below because its fluorescence signal was extremely high for this yellow/red filter set; intermediate bands in this ladder are not strongly fluorescent in either channel. Although both bright fluorescent bands give background in adjacent lanes, this did not significantly impact measuring the intensity of either PerM^n1^-mNG band of interest. (**B**) The same gel as above was imaged with a blue/green filter set; in these extraction conditions used to quantify PerM^n1^-mNG degradation, full-length mTq2-FtsB constructs are not solubilized; instead, degradation products with mTurquoise2 fluorescence are observed near the expected 27 kD molecular weight of mTurquoise2.

## References

[1] Houben, R.M.G.J. & Dodd, P. J. The Global Burden of Latent Tuberculosis Infection: A Re-estimation Using Mathematical Modelling. PLOS Medicine 13, e1002152 (2016). 10.1371/journal.pmed.1002152.

[2] WHO. Latent tuberculosis infection: Updated and consolidated guidelines for programmatic management. Tech. Rep. (2018).

[3] Dale, K. D. et al. Quantifying the rates of late reactivation tuberculosis: A systematic review. The Lancet Infectious Diseases 21, e303–e317 (2021). 10.1016/S1473-3099(20)30728-3.

[4] Assefa, D. G. et al. Efficacy and safety of different regimens in the treatment of patients with latent tuberculosis infection: A systematic review and network meta-analysis of randomized controlled trials. Archives of Public Health 81, 82 (2023). 10.1186/s13690-023-01098-z.

[5] Li, T.-L. et al. Acquired Resistance to Isoniazid During Isoniazid Monotherapy in a Subject with Latent Infection Following Household Rifampicin-Resistant Tuberculosis Contact: A Case Report. Infect Drug Resist 14, 1505–1509 (2021). 10.2147/IDR.S304799.

[6] Dartois, V. A. & Rubin, E. J. Anti-tuberculosis treatment strategies and drug development: Challenges and priorities. Nat Rev Microbiol 20, 685–701 (2022). https://doi.org/10/gqbtrn.

[7] Vandal, O. H., Pierini, L. M., Schnappinger, D., Nathan, C. F. & Ehrt, S. A membrane protein preserves intrabacterial pH in intraphagosomal Mycobacterium tuberculosis. Nat Med 14, 849–854 (2008). https://doi.org/10/drbt2b.

[8] Goodsmith, N. et al. Disruption of an M. tuberculosis Membrane Protein Causes a Magnesium-dependent Cell Division Defect and Failure to Persist in Mice. PLOS Pathogens 11, e1004645 (2015). https://doi.org/10/f67pgq.

[9] Wang, R. et al. Persistent Mycobacterium tuberculosis infection in mice requires PerM for successful cell division. eLife 8, e49570 (2019). 10.7554/eLife.49570. 10. | |

[10] Meade, R. K. et al. Genome-wide screen identifies host loci that modulate Mycobacterium tuberculosis fitness in immunodivergent mice. G3 Genes Genomes Genetics 13, jkad147 (2023). https://doi.org/10/gtm6dn.

[11] Paulussen, F. M. et al. Covalent Proteomimetic Inhibitor of the Bacterial FtsQB Divisome Complex. Journal of the American Chemical Society 144, 15303–15313 (2022). https://doi.org/10/gtm6dp.

[12] Wu, K. J. et al. Characterization of Conserved and Novel Septal Factors in Mycobacterium smegmatis. J Bacteriol 200, e00649–17 (2018). https://doi.org/10/gtm6dr.

[13] Grāve, K., Bennett, M. D. & Högbom, M. High-throughput strategy for identification of Mycobacterium tuberculosis membrane protein expression conditions using folding reporter GFP. Protein Expr Purif 198, 106132 (2022). 10.1016/j.pep.2022.106132.

[14] Korepanova, A. et al. Cloning and expression of multiple integral membrane proteins from Mycobacterium tuberculosis in Escherichia coli. Protein Sci 14, 148–158 (2005). https://doi.org/2023101017072700228.

[15] Baek, M. et al. Accurate prediction of protein structures and interactions using a three-track neural network. Science 373, 871–876 (2021). https://doi.org/10/gk7nhq.

[16] Evans, R. et al. Protein complex prediction with AlphaFold-Multimer. bioRxiv 2021.10.04.463034 (2022). 10.1101/2021.10.04.463034.

[17] Attaibi, M. & den Blaauwen, T. An Updated Model of the Divisome: Regulation of the Septal Peptido-glycan Synthesis Machinery by the Divisome. Int J Mol Sci 23, 3537 (2022). 10.3390/ijms23073537.

[18] Craven, S. J., Condon, S. G. F. & Senes, A. A model of the interactions between the FtsQLB and the FtsWI peptidoglycan synthase complex in bacterial cell division. bioRxiv 2022.10.30.514410 (2022). 10.1101/2022.10.30.514410.

[19] Käshammer, L. et al. Cryo-EM structure of the bacterial divisome core complex and antibiotic target FtsWIQBL. Nat Microbiol 1–11 (2023). https://doi.org/10/gtm6dm.

[20] Britton, B. M. et al. Conformational changes in the essential E. coli septal cell wall synthesis complex suggest an activation mechanism. Nat Commun 14, 4585 (2023). https://doi.org/10/gtm4j5.

[21] Park, K.-T., Park, D. J., Pichoff, S., Du, S. & Lutkenhaus, J. The essential domain of FtsN triggers cell division by promoting interaction between FtsL and FtsI. bioRxiv 2023.05.12.540521 (2023). https://doi.org/10/mmwf.

[22] Bryant, P., Pozzati, G. & Elofsson, A. Improved prediction of protein-protein interactions using AlphaFold2. Nat Commun 13, 1265 (2022). 10.1038/s41467-022-28865-w.

[23] Varadi, M. et al. AlphaFold Protein Structure Database: Massively expanding the structural coverage of protein-sequence space with high-accuracy models. Nucleic Acids Research 50, D439–D444 (2022). 10.1093/nar/gkab1061.

[24] Shaner, N. C. et al. A bright monomeric green fluorescent protein derived from Branchiostoma lanceolatum. Nat Methods 10, 407–409 (2013). https://doi.org/10/f24459.

[25] Goedhart, J. et al. Structure-guided evolution of cyan fluorescent proteins towards a quantum yield of 93%. Nat Commun 3, 751 (2012). 10.1038/ncomms1738.

[26] Lee, P. A., Tullman-Ercek, D. & Georgiou, G. The Bacterial Twin-Arginine Translocation Pathway. Annu Rev Microbiol 60, 373–395 (2006). 10.1146/annurev.micro.60.080805.142212.

[27] Kiefer, D., Hu, X., Dalbey, R. & Kuhn, A. Negatively charged amino acid residues play an active role in orienting the Sec-independent Pf3 coat protein in the Escherichia coli inner membrane. The EMBO Journal 16, 2197–2204 (1997). 10.1093/emboj/16.9.2197.

[28] Lucena, D., Mauri, M., Schmidt, F., Eckhardt, B. & Graumann, P. L. Microdomain formation is a general property of bacterial membrane proteins and induces heterogeneity of diffusion patterns. BMC Biol 16, 97 (2018). 10.1186/s12915-018-0561-0.

[29] Heckert, A., Dahal, L., Tjian, R. & Darzacq, X. Recovering mixtures of fast-diffusing states from short single-particle trajectories. eLife 11, e70169 (2022). 10.7554/eLife.70169.

[30] Hansen, A. S. et al. Robust model-based analysis of single-particle tracking experiments with Spot-On. eLife 7, e33125 (2018). 10.7554/eLife.33125.

[31] Wang, R. & Ehrt, S. Rv0954 Is a Member of the Mycobacterial Cell Division Complex. Frontiers in Microbiology 12 (2021). https://doi.org/10/gm8vsc.

[32] Baranowski, C., Rego, E. H. & Rubin, E. J. The Dream of a Mycobacterium. Microbiology Spectrum 7, 10.1128/microbiolspec.gpp3–0008–2018 (2019). https://doi.org/10/gtm6dj.

[33] Shlosman, I. et al. Allosteric activation of cell wall synthesis during bacterial growth. Nat Commun 14, 3439 (2023). 10.1038/s41467-023-39037-9.

[34] Mirdita, M. et al. ColabFold: Making protein folding accessible to all. Nat Methods 19, 679–682 (2022). 10.1038/s41592-022-01488-1.

[35] Steinegger, M. & Söding, J. MMseqs2 enables sensitive protein sequence searching for the analysis of massive data sets. Nat Biotechnol 35, 1026–1028 (2017). https://doi.org/10/ggctnw.

[36] DeLano, W. L. Pymol: An open-source molecular graphics tool. CCP4 Newsletter on protein crystallography 40, 82–92 (2002).

[37] Sievers, F. et al. Fast, scalable generation of high-quality protein multiple sequence alignments using Clustal Omega. Mol Syst Biol 7, 539 (2011). https://doi.org/10/bmv826.

[38] Kunzmann, P. & Hamacher, K. Biotite: A unifying open source computational biology framework in Python. BMC Bioinformatics 19, 346 (2018). 10.1186/s12859-018-2367-z.

[39] Hallgren, J. et al. DeepTMHMM predicts alpha and beta transmembrane proteins using deep neural networks. bioRxiv 2022.04.08.487609 (2022). 10.1101/2022.04.08.487609.

[40] Tareen, A. & Kinney, J. B. Logomaker: Beautiful sequence logos in Python. Bioinformatics 36, 2272–2274 (2020). 10.1093/bioinformatics/btz921.

[41] Wu, E. L. et al. CHARMM-GUI Membrane Builder toward realistic biological membrane simulations. Journal of Computational Chemistry 35, 1997–2004 (2014). https://doi.org/10/f6kgzz.

[42] Huang, J. et al. CHARMM36m: An improved force field for folded and intrinsically disordered proteins. Nat Methods 14, 71–73 (2017). https://doi.org/10/gfxkj3.

[43] Eastman, P. et al. OpenMM 7: Rapid development of high performance algorithms for molecular dynamics. PLOS Computational Biology 13, e1005659 (2017). https://doi.org/10/gbppkv.

[44] Brown, T., Chavent, M. & Im, W. Molecular Modeling and Simulation of the Mycobacterial Cell Envelope: From Individual Components to Cell Envelope Assemblies. J. Phys. Chem. B 127, 10941–10949 (2023). 10.1021/acs.jpcb.3c06136.

[45] Suh, D. et al. CHARMM-GUI Enhanced Sampler for various collective variables and enhanced sampling methods. Protein Sci 31, e4446 (2022). 10.1002/pro.4446.

[46] Miao, Y., Nichols, S. E., Gasper, P. M., Metzger, V. T. & McCammon, J. A. Activation and dynamic network of the M2 muscarinic receptor. Proceedings of the National Academy of Sciences 110, 10982–10987 (2013). 10.1073/pnas.1309755110.

[47] Gao, Y. et al. CHARMM-GUI Supports Hydrogen Mass Repartitioning and Different Protonation States of Phosphates in Lipopolysaccharides. Journal of Chemical Information and Modeling 61, 831–839 (2021). 10.1021/acs.jcim.0c01360.

[48] Gowers, R. J. et al. MDAnalysis: A Python Package for the Rapid Analysis of Molecular Dynamics Simulations. Proceedings of the 15th Python in Science Conference 98–105 (2016). 10.25080/Majora-629e541a-00e.

[49] Michaud-Agrawal, N., Denning, E. J., Woolf, T. B. & Beckstein, O. MDAnalysis: A toolkit for the analysis of molecular dynamics simulations. J Comput Chem 32, 2319–2327 (2011). 10.1002/jcc.21787.

[50] Zhang, M. et al. Rational design of true monomeric and bright photoactivatable fluorescent proteins. Nat Methods 9, 727–729 (2012). 10.1038/nmeth.2021.

[51] Hensel, Z. A plasmid-based Escherichia coli gene expression system with cell-to-cell variation below the extrinsic noise limit. PLOS ONE 12, e0187259 (2017). https://doi.org/10/gcg8pf.

[52] Silva, J. P. N., Lopes, S. V., Grilo, D. J. & Hensel, Z. Plasmids for Independently Tunable, Low-Noise Expression of Two Genes. mSphere 4 (2019). https://doi.org/10/gf3vhf.

[53] Neidhardt, F. C., Bloch, P. L. & Smith, D. F. Culture Medium for Enterobacteria. J Bacteriol 119, 736–747 (1974).

[54] Schindelin, J. et al. Fiji: An open-source platform for biological-image analysis. Nature methods 9, 676–82 (2012). https://doi.org/10/f34d7c.

[55] Cutler, K. J. et al. Omnipose: A high-precision morphology-independent solution for bacterial cell segmentation. Nat Methods 19, 1438–1448 (2022). 10.1038/s41592-022-01639-4.

[56] Sanial, M. et al. Direct observation of fluorescent proteins in gels: A rapid cost-efficient, and quantitative alternative to immunoblotting. bioRxiv 2024.05.31.594679 (2024). 10.1101/2024.05.31.594679.

[57] Tinevez, J.-Y. et al. TrackMate: An open and extensible platform for single-particle tracking. Methods 115, 80–90 (2017). 10.1016/j.ymeth.2016.09.016.

